# Conservation assessment of human splice site annotation based on a 470-genome alignment

**DOI:** 10.1101/2023.12.01.569581

**Authors:** Ilia Minkin, Steven L. Salzberg

## Abstract

Despite many improvements over the years, the annotation of the human genome remains imperfect. The use of evolutionarily conserved sequences provides a strategy for selecting a high-confidence subset of the annotation. Using the latest whole genome alignment, we found that splice sites from protein-coding genes in the high-quality MANE annotation are consistently conserved across more than 350 species. We also studied splice sites from the RefSeq, GENCODE, and CHESS databases not present in MANE. In addition, we analyzed the completeness of the alignment with respect to the human genome annotations and described a method that would allow us to fix up to 50% of the missing alignments of the protein-coding exons. We trained a logistic regression classifier to distinguish between the conservation exhibited by sites from MANE versus sites chosen randomly from neutrally evolving sequences. We found that splice sites classified by our model as well-supported have lower SNP rates and better transcriptomic evidence. We then computed a subset of transcripts using only “well-supported” splice sites or ones from MANE. This subset is enriched in high-confidence transcripts of the major gene catalogs that appear to be under purifying selection and are more likely to be correct and functionally relevant.

**Graphical abstract:** 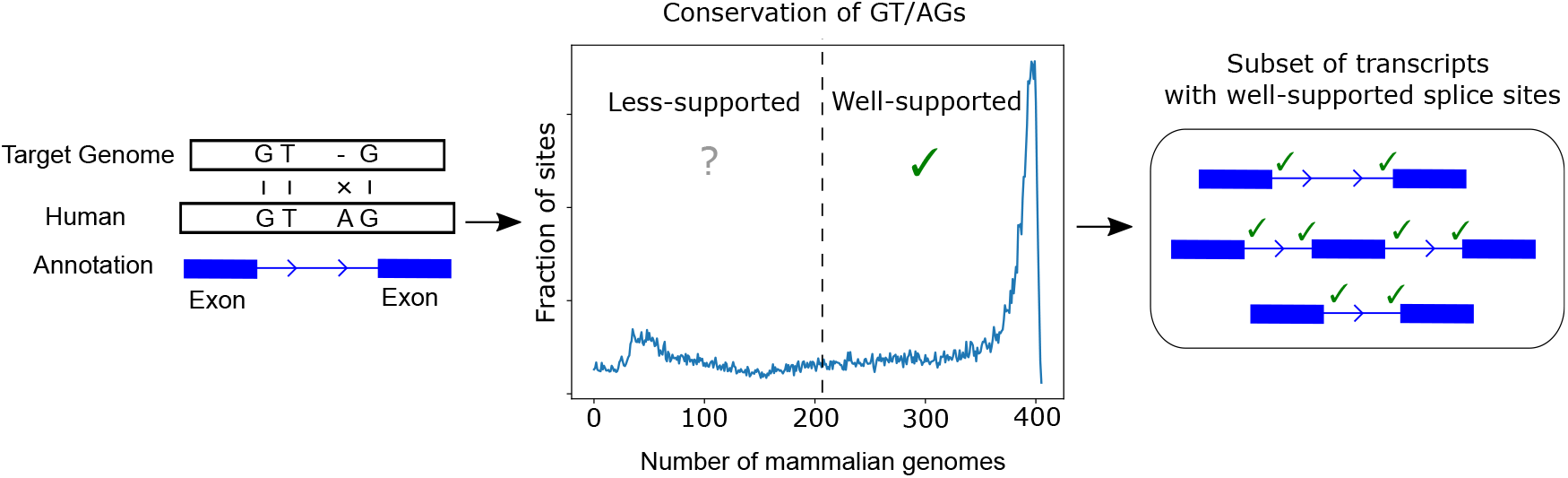

## Introduction

The annotation of the human genome is a fundamental resource for a broad range of biomedical research and clinical applications. However, more than two decades after the initial publication of the genome itself, the scientific community have not reached a point where a consensus genome annotation is available [1]. For example, one consequence is that the leading gene annotation databases for the human reference genome often disagree even on basic statistics such as the number of protein-coding genes [39]. This is due to a variety of reasons, including the imperfect technologies used to assemble RNA transcripts and the noise inherent in the transcription process itself [44, 56, 7].

One of the challenging aspects of constructing a genome annotation is correctly determining the positions of introns inside the genes. The existence of introns and the mechanism of alternative splicing, first proposed by Gilbert [18], are critical for the functioning of cells. At the same time, the evolutionary origin of introns has been the subject of a scientific debate for decades [12, 32, 47, 25].

A recent effort to address the challenge of the discrepancy between different human annotations resulted in the creation of a limited, high-quality gene annotation database called MANE [36]. This annotation was intended to include a single representative transcript for each protein-coding gene that has identical exon and intron structures in both RefSeq and GENCODE, two of the leading human annotation databases. The transcripts in MANE are chosen based on criteria that include expression levels and evolutionary conservation, which is a strong predictor of biological function. A similar project called APPRIIS [46] provides a single transcript for every protein-coding gene based on human genetics data, protein evidence, and cross-species conservation; APPRIS contains annotations for the human as well as a few other reference species. These approaches yielded a subset of the human transcriptome under strong purifying selection. These and other studies of the evolutionary consistency of the human genome annotation [14] were mostly focused on the sequences of the protein-coding exons rather than splice site motifs.

In this study, we address the question of the conservation of splice sites in major gene catalogs, both across multiple species and population levels. First, we analyzed the completeness of the alignment containing 470 mammalian species recently published by the UCSC Genome Browser team [45] with respect to the annotation of the human exons; we restricted this alignment to 405 species due to sequence availability reasons. As we observed alignments of many exons to be missing, we came up with a method to fix the missing alignment, recovering up to 50% of the missing exon/genomes pairs. Second, we observed that the canonical dinucleotides GT/AG that flank introns are very highly conserved in protein-coding genes in MANE, genomes with most of them being intact in more than 350 species. We then investigated the patterns of conservation among splice sites that are not in MANE but that are present in one or more of the leading gene catalogs RefSeq, GENCODE, and CHESS. We found that while many of those splice sites closely follow the pattern of conservation found in MANE, others resemble randomly generated sites from neutrally evolving sequences.

To compare the properties of these two groups of splice sites, we developed a logistic regression model that classifies splice sites as either well-supported or less-supported. The model relies on a comparison of conservation patterns of splice sites from MANE to neutrally evolving sequences. As we detail below, we found that sites predicted as well-supported by our classifier have lower rates of single nucleotide polymorphisms (SNPs) in the human population, are enriched in clinically relevant polymorphisms, and have better transcriptomic support. We then obtained a subset of transcripts from each major gene catalog for which all splice sites were either classified as well-supported by our model or included in a transcript from MANE. These transcripts appear to be under strong purifying selection and are more likely to be functional and clinically relevant.

## Methods

### Realignment of missing exon/genome pairs

Before investigating the conservation of the splice sites, we performed a procedure to fix the gaps in the alignment that might affect the results. First, we found human exons and particular genomes such that the exon is not aligned anywhere in that target genome. We then tried to realign these exons using the synteny information. The intuition is that if a human exon is not aligned to another genome, but down- and up-stream exons are mapped to the same sequence in that target genome, then we can try to place the missing exon in between two of its neighbors in the target genome. Below we give a more detailed description of the method.

We are given a collection of genomes *G* = {*g*_1_, …, *g*_*m*_}, where each genome is a string 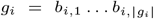 over the *nucleotides* of the DNA alphabet {*A, C, G, T* }, where |*g*_*i*_| is the length of the *i*-th genome. The genome *g*_1_ is called the *reference*, and any non-reference genome *g*_*t*_, *t >* 1 is called a *target* genome. For the reference genome, we are given an *exon annotation* represented as a set of segments *E* = {(*x*_1_, *y*_1_), …, (*x*_*t*_, *y*_*t*_)}, 1 ≤ *x*_*i*_ *< y*_*i*_ ≤ |*g*_1_|.

To find the corresponding sequence of each exon of the reference in another species, we use a whole-genome alignment of *m* species. Formally, we define an alignment function *w*(*k, g*_*t*_) that maps each position *k* of the reference genome to its homologous position in target genome *g*_*t*_ included in the alignment if such position exists, otherwise, *w*(*k, g*_*t*_) = −1.

We say that an exon *e* = (*x, y*) ∈ *E* is *unaligned* in target genome *g*_*t*_ if *w*(*k, g*_*t*_) = −1 for all *x* ≤ *k* ≤ *y*; otherwise we call the exon *aligned*. We define the set of all aligned positions of exon *e* as *A*(*e*) = {*w*(*k, g*_*t*_)|*x* ≤ *k* ≤ *y, w*(*k, g*_*t*_) ≠ −1}. We call an exon *e*_*i*_ *syntenic* in genome *g*_*t*_ if there are two other exons *e*_*a*_ = (*x*_*a*_, *y*_*a*_), *e*_*b*_ = (*x*_*b*_, *y*_*b*_), *y*_*a*_ *< x < y < x*_*b*_ that are aligned in *g*_*t*_.

We use the fact that unaligned, but syntenic exons have other neighboring exons mapped to the target genome to get a hint of where the alignment of the said exons could be. Let *e* be such an unaligned exon. Then the target segment *u* is defined as *u* = (max(*A*(*e*_*a*_)) + 1, min(*A*(*e*_*b*_)) − 1). We use edlib library [55] to find the best alignment of *e* to the range *u* in the target genome *g*_*t*_ to get the alignment function *w*_*e*_(*k, g*_*t*_) by aligning the specific exon *e*. After trying to realign all such exons *e*, we merge the resulting alignments *w*_*e*_(*k, g*_*t*_) with the original alignment *w*(*k, g*_*t*_); we define the resulting function as *w*^′^(*k, g*_*t*_). Figure 1a illustrates the above definitions.

**Fig. 1.**
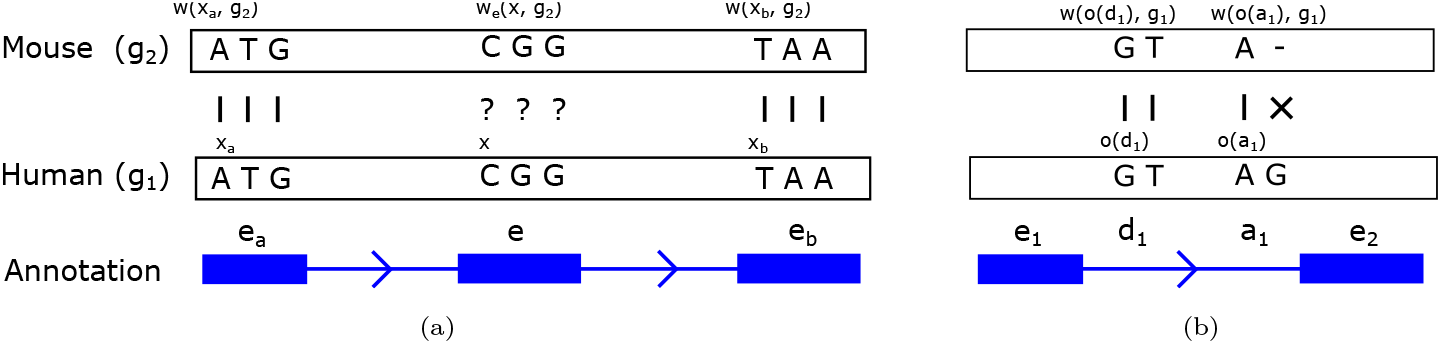
An example demonstrating the definitions from the Methods section. Panel (a) shows an alignment of exons from the reference genome *g*_1_ (Human) to a target genome *g*_2_ (Mouse) using a whole-genome alignment. Blue boxes represent positions of exons *e*_*a*_, *e*, and *e*_*b*_ from the annotation of the Human genome, and the arrowed lines are introns; splice sites are not shown for visual clarity. In this example, exons *e*_*a*_ and *e*_*b*_ are *aligned*, with vertical dashes indicating the alignment between nucleotides of the different genomes. The first positions of the corresponding exons *x*_*a*_ and *x*_*b*_ are aligned to their counterparts in the target genomes, *w*(*x*_*a*_, *g*_2_) and *w*(*x*_*b*_, *g*_2_). The exon *e* is unaligned since all its positions are missing in the alignment, as indicated by question marks, *w*(*x, g*_2_) = *w*(*x*+1, *g*_2_) = *w*(*x*+2, *g*_2_) = *−*1. At the same time, the exon *e* is syntenic, since its neighboring exons are aligned, and we can reasonably hypothesize that *e* can be aligned to the segment between the alignments of *e*_*a*_ and *e*_*b*_, or *w*(*x*_*a*_, *g*_2_) *< w*_*e*_(*x, g*_2_) *< w*(*x*_*b*_, *g*_2_). Panel (b) shows an example of splice sites annotation: *d*_1_ denote the position *o*(*d*_1_) of the first of the canonical dinucleotides of the donor splice site, and *a*_1_ denote the position of the first of the canonical dinucleotides *o*(*a*_1_) of the acceptor site. The donor site *d*_1_ of the human reference genome *g*_1_ has both its canonical dinucleotides intact in the target mouse genome, *g*_2_. However, this is not true for the acceptor site *a*_1_ mutated in mouse. In this example, the values of the conservation function for these two splice sites are *C*(*d*_1_, 0, 2) = *C*(*d*_1_, 1, 2) = 1, *C*(*a*_1_, 0, 2) = 0, and *C*(*a*_1_, 1, 2) = 1.

We apply several filtering steps along the process. To reduce the computation load, we only consider target segments *u* with a length less than a predefined threshold, which we set to 100,000 in our experiments. We also used the following criteria to filter out potentially spurious alignments. Let *E*_*t*_ ⊆ *E* be a subset of exons aligned in the genome *g*_*t*_. For an exon *e* = (*x, y*) ∈ *E*_*t*_ we define its alignment score 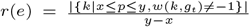, or as a fraction of the position of the exon *e* that are aligned in *g*_*t*_. Let *R*_*t*_ = {*r*(*e*)|*e* ∈ *E*_*t*_} be the set of the scores of all aligned exons with respect to the genome *g*_*t*_ in the original alignment. We only accept a realignment *w*_*e*_(*k, g*_*t*_) of exon *e* = (*x, y*) if the alignment score 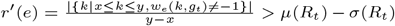, where *μ* and *σ* are arithmetic mean and standard deviation correspondingly.

### Splice site classification

Our method for classifying splice sites is based on a logistic regression model designed to predict the probability of a splice site having a MANE-like conservation pattern (well-supported), or a conservation pattern similar to a neutrally evolving sequence (less-supported). One of the primary features used by the regression model is the number of species in which the canonical dinucleotides are conserved, computed from a large multiple genome alignment. In addition, it takes into account an array of positions surrounding a splice site, as they appear to have similar conservation properties. Having such a classification method in addition to the number of species in which the canonical dinucleotides are conserved is necessary because the number itself is not informative without a baseline representing neutrally evolving sequences to compare against. The training data includes randomly chosen sites from intronic sequences as negative examples and the whole MANE dataset as positive examples. Below, we give the necessary initial definitions and describe the model.

We are given a *splice site annotation* for the reference genome, represented as two sets, donor sites *D* = {*d*_1_, …, *d*_|*D*|_}, and acceptor sites *A* = {*a*_1_, …, *a*_|*A*|_}. The *origin* of the site is the position of the nucleotide of the first of the canonical dinucleotides, which we designate as *o*(*s*) where *s* is either a donor or splice site. Thus for most donor sites 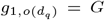 and 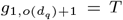 and for most acceptor sites 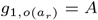 and 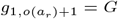.

To find the corresponding sequence of each splice site of the reference in another species, we use a whole-genome alignment of *m* species represented as the alignment function *w*^′^(*p, g*_*t*_) that maps a position *p* of the reference genome to a target genome *g*_*t*_. We also define the conservation function *C*(*s, ℓ, t*) as follows: it takes the value of 1 if the nucleotide with the shift *ℓ* of splice site *s* matches its homologous nucleotide in the genome *t* and 0 otherwise: *C*(*s, ℓ, t*) = *I*[*b*_1,*o*(*s*)+*ℓ*_ = *b*_*t*,*w*(*o*(*s*)+*ℓ*,*t*)_]. Figure 1b shows an example of mapping splice site sequence using whole-genome alignment and computation of the alignment and conservation function.

Our model consists of two types of variables to classify a splice site as well-supported or less-supported: (1) number of species in which the canonical dinucleotides are conserved *jointly* (2) number of species in which each nucleotide 30 position down- and up-stream of the canonical dinucleotide is conserved in, one variable per each position. This way, the log-odds of an acceptor site *a*_*r*_ being well-supported are defined as:

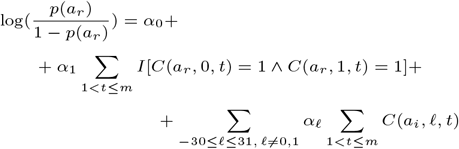

Where *α*_0_ is the interceptor term, *α*_1_ corresponds to the conservation of canonical dinucleotides, and *α*_*ℓ*_ is the coefficient corresponding to the conservation of the position with the shift *ℓ* of the splice site. The log-odds of a donor splice being well-supported are defined analogously. In addition, we evaluated a model taking into account the conservation of the canonical dinucleotides only to evaluate the contribution of the rest of the positions in the splicing motif. This way, we define the log-odds with the respect to the probability *p*_0_ of an acceptor splice site *a*_*r*_ being well-supported as:

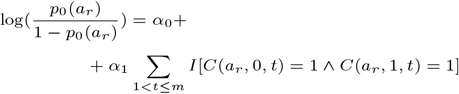

### Data and software acquisition

We analyzed the following gene catalogs: GENCODE [15] version 45, RefSeq [37] release 110, CHESS 3 [59] v.3.1.0, and MANE [36] Select v1.3. We note that GENCODE, RefSeq, and CHESS 3 all contain every gene and transcript in MANE, which was created by GENCODE and RefSeq scientists with the goal of providing a single high-confidence transcript for every human protein-coding gene. To take into account this confounding factor and observe the differences between annotations more clearly, we removed the MANE splice sites from each of the other catalogs and created reduced versions that we designate as GENCODE^∗^, RefSeq^∗^, and CHESS 3^∗^ respectively. This procedure only affected protein-coding genes, because MANE does not currently contain non-coding genes or other types of annotation.

We only included protein-coding and lncRNA genes in our analyses; however, the way these catalogs define gene types slightly differs. GENCODE and MANE, we used the attribute “transcript type” of a transcript to infer its type; for CHESS 3, we used the attribute “gene type” of a transcript for this purpose. For RefSeq, we consider a transcript to be protein-coding if its corresponding gene was assigned “protein coding” to its “gene biotype” attribute, and the transcript was assigned “mRNA” to its “transcript biotype” attribute. For lncRNAs from RefSeq, we consider a transcript to be lncRNA if its corresponding gene was assigned “lncRNA” to its “gene biotype” attribute, and the transcript was assigned “lnc RNA” to its “transcript biotype” attribute. We note that for protein-coding genes we considered all introns from the mentioned annotations, which includes the ones located in untranslated regions (UTRs).

In addition, we created a false gene annotation, intended to capture a baseline of neutrally evolving sequences; we refer to this dataset as “Random.” This annotation consists of 180,000 randomly generated transcripts located within introns of genes of MANE outside of splicing motifs. Each transcript consists of two short exons separated by an intron, yielding 180,000 distinct donor and acceptor sites.

We used a 470-species alignment available at the UCSC Genome Browser website [26] generated using MultiZ whole-genome aligner [5]. However, we had to restrict this alignment to 405 species: in order to implement our exon realignment procedure, we also had to download the sequences of the genomes themselves, and 65 of these genomes were unavailable for download; the full list of 405 genomes is available in online documentation. In addition, we excluded any genes located on the “patches” sequences of GRCh38 [50] if such sequences were not included in the original alignment produced by UCSC; we also excluded single exon transcripts. We also excluded chromosome Y from our analysis due to the reasons listed in the Results section.

For fitting coefficients of the regression equations above, we used the logistic regression module from SciKit [38]. To implement the alignment function *w*(*k, g*_*t*_), we utilized the AlignIO library from BioPython module [9] version 1.79.

## Results

### Assessing completeness of the alignment

Any conclusions about the conservation of splice sites drawn from the alignment analysis depend on its completeness. To assess it, we calculated the following statistics that reflect how many exons from human gene catalogs are mapped to the other genomes. Let *E* be the set of protein-coding and lncRNA exons from the three gene catalogs under consideration and *G* be the genomes used in the whole-genome alignment.

We define a variable *W* (*e, g*), *e* ∈ *E, g* ∈ *G* that indicates whether exon *e* is aligned with the genome *g* as follows. We assign *W* (*e, g*) = 1 if at least one position of *e* is aligned somewhere in the genome *g*, and *W* (*e, g*) = 0 otherwise. To present the summary of the alignment of the exons, we calculate the sums *s*_*m*_ = ∑ (1 − *W* (*e, g*)) and *s*_*a*_ = ∑*W* (*e, g*), showing how many exon/genome pairs are missing and present in the alignment correspondingly.

Table 1, columns 2-5 represent these statistics *s*_*a*_ and *s*_*m*_ broken down by chromosome and gene type. For autosomal chromosomes, we observe that up to 13% exon/genome pairs are missing for protein-coding genes, and up to 50% of such pairs are missing for lncRNAs. Sex chromosome Y is an obvious outlier since more than 35% of protein-coding and 72% of lncRNA exon/genome pairs are missing in the alignment.

**Table 1.** The number of aligned protein-coding and lncRNA exons/genomes in the original alignment of 470 mammals restricted to 405 species (columns 2-3, defined as *s*_*a*_ in the main text), missing in the alignment (columns 4-5, defined as *s*_*m*_ in the main text), and recovered using our synteny-based realignment procedure (columns 6-7, defined *s*_*r*_ in the main text). We computed these numbers for the union of the set of exons of all gene annotations under consideration: GENCODE, RefSeq, CHESS 3, and MANE.

|  | Gene type | Aligned exons/genomes |  | Missing exons/genomes |  | Recovered exons/genomes |  |
| --- | --- | --- | --- | --- | --- | --- | --- |
|  |  | Coding | lncRNA | Coding | lncRNA | Coding | lncRNA |
| Chromosome | 1 | 13,524,670 | 4,943,971 | 1,173,995 | 1,848,284 | 558,122 | 1,016,754 |
|  | 2 | 10,200,614 | 5,196,973 | 764,761 | 1,798,592 | 426,570 | 1,057,443 |
|  | 3 | 8,517,408 | 3,747,144 | 569,172 | 1,263,921 | 331,051 | 743,381 |
|  | 4 | 5,524,742 | 2,887,223 | 504,088 | 1,325,587 | 274,431 | 754,267 |
|  | 5 | 6,143,565 | 3,287,189 | 404,475 | 1,260,556 | 230,366 | 684,250 |
|  | 6 | 6,669,798 | 3,437,254 | 544,467 | 1,290,311 | 295,057 | 756,804 |
|  | 7 | 6,478,768 | 2,808,452 | 589,292 | 1,288,933 | 312,680 | 666,229 |
|  | 8 | 4,886,473 | 2,744,951 | 478,967 | 1,302,619 | 259,518 | 684,356 |
|  | 9 | 5,271,324 | 2,320,005 | 404,346 | 993,705 | 215,712 | 435,116 |
|  | 10 | 5,399,044 | 2,491,811 | 456,446 | 989,569 | 257,332 | 561,440 |
|  | 11 | 8,459,650 | 2,620,330 | 653,255 | 844,445 | 336,269 | 501,208 |
|  | 12 | 8,110,205 | 2,751,895 | 583,525 | 1,148,660 | 332,270 | 663,752 |
|  | 13 | 2,334,841 | 1,663,448 | 217,469 | 794,497 | 131,064 | 418,293 |
|  | 14 | 4,676,912 | 2,022,926 | 304,993 | 688,144 | 187,024 | 392,903 |
|  | 15 | 4,948,431 | 2,327,369 | 341,679 | 834,871 | 185,590 | 425,251 |
|  | 16 | 6,284,584 | 2,227,608 | 602,846 | 824,067 | 320,788 | 454,321 |
|  | 17 | 8,710,715 | 2,248,362 | 685,690 | 868,518 | 341,519 | 441,792 |
|  | 18 | 2,366,477 | 1,501,463 | 191,098 | 606,967 | 112,751 | 366,562 |
|  | 19 | 8,336,039 | 1,275,528 | 1,333,336 | 1,222,917 | 620,313 | 483,427 |
|  | 20 | 3,371,434 | 1,546,869 | 259,391 | 636,486 | 150,521 | 351,570 |
|  | 21 | 1,299,495 | 1,000,286 | 165,795 | 561,799 | 93,259 | 256,060 |
|  | 22 | 2,908,246 | 1,043,948 | 309,884 | 516,517 | 160,708 | 228,741 |
|  | X | 4,774,991 | 1,068,688 | 597,739 | 648,917 | 200,967 | 191,003 |
|  | Y | 236,212 | 130,554 | 147,728 | 350,991 | 20,636 | 32,383 |

Since we observed a significant amount of exon/genome pairs not being aligned, we developed a strategy to recover them using the synteny information. The description of this step can be found in the subsection “Realignment of missing exon/genome pairs” of the “Methods.” We define this quantity as *s*_*r*_ = ∑ *W*_*r*_(*e, g*), for *e* ∈ *E, g* ∈ *G* such that *W* (*e, g*) = 0, where *W*_*r*_(*e, g*) = 1 if at least on position of the exon *e* is aligned somewhere in the genome *g* by the extended alignment function described in the subsection “Realignment of missing exon/genome pairs” of the “Methods.” Table 1, columns 6-7 show the number of exon/genome pairs recovered by our method: for autosomal chromosomes, we recovered up to 60% of exon/genome pairs for both protein-coding genes and lncRNAs. In contrast, we recovered 33-29% of such pairs for chromosome X and only 9-13% of the exon/genome pairs for chromosome Y. This can be explained by the fact that many genomes are missing the assembled Y chromosome and its challenging structure, which is comprised of rich families of repeated sequences. Hence, we decided to exclude chromosome Y from our analysis. In addition to realigning the exons from the real datasets, we realigned the exons from the “Random” dataset to have a realistic baseline. Supplementary Table S1 shows these numbers that are similar to the real gene annotations: we were able to recover 58-66% percent of exon/genome pairs.

### Exploratory data analysis of splice site conservation

First, we evaluated the evolutionary conservation of splice sites from four different human genome annotation databases: GENCODE(*), RefSeq(*), CHESS 3(*), and MANE Select. Table 2 shows the numbers of donor and acceptor sites in each dataset, as well as the total number of transcripts. For every donor and acceptor splice site in the databases, we computed how many species preserve the consensus dinucleotides (GT and AG) that appear at the beginning and end of most introns. Figure 2 shows the pattern of conservation across species for each of these sets of donor and acceptor sites. First, we note that splice sites from protein-coding genes in MANE yield a plot that is clearly distinct from the other gene catalogs: most of the sites from MANE are conserved in *>* 350 species. Second, protein-coding splice sites from the other datasets (after removing the MANE splice sites) seem to fall into two distinct categories: (1) MANE-like, and (2) neutral-like conservation. We also noted a small peak for splice sites from protein-coding genes of RefSeq^∗^ at around 330 species: most of these splice sites come from “patches” sequences to the hg38 reference genome that are absent in the other annotations. In contrast, lncRNAs from all datasets have very similar distributions that closely follow the conservation pattern of Random 180,000 180,000 - - 180,000 - random sites. Both donor and acceptor splice show similar patterns of conservation. We note that randomly generated sites along with lncRNAs and some sites from coding genes exhibit several peaks in conservation in fewer than 50 species.

**Table 2.**
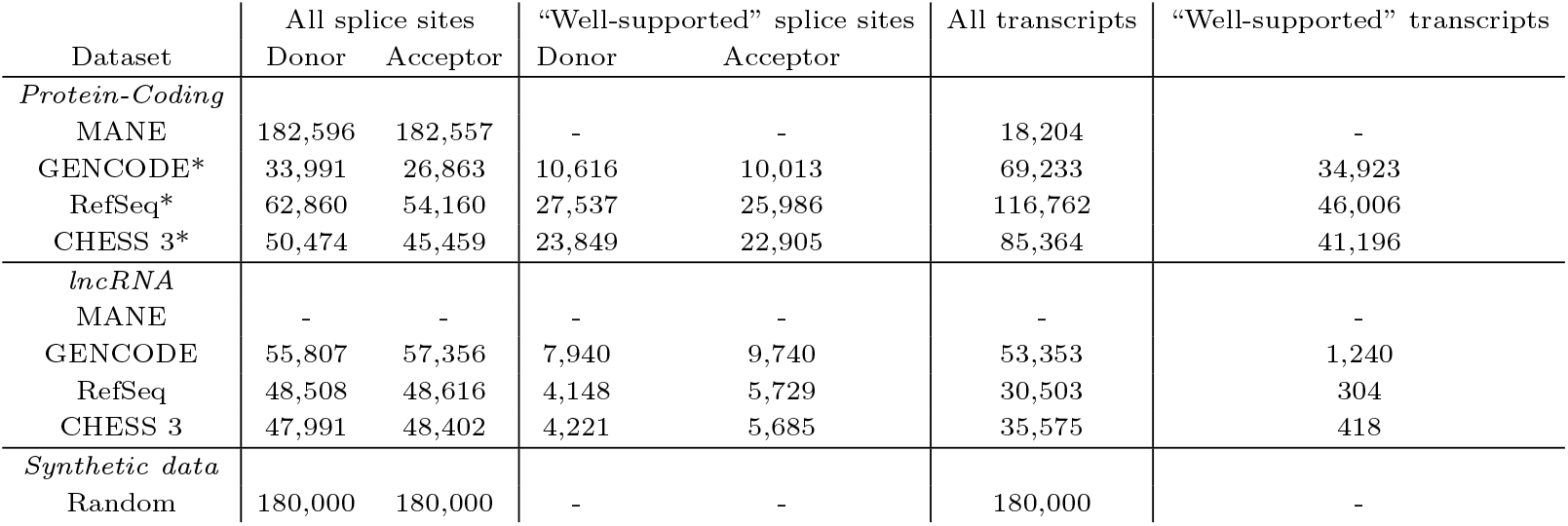
Summary statistics of splice site conservation analysis. The second and the third columns represent the total number of donor and acceptor sites in each dataset and the third and fourth columns show the number of donor and acceptor splice sites classified as “well-supported” by our model. The last two columns indicate the total number of transcripts in each dataset and the number of transcripts that have all splice sites either from MANE dataset or classified as “well-supported.” We only considered transcripts with at least one intron present. Dashes indicate that transcripts and splice sites of a certain type were not available in a dataset.

**Fig. 2.**
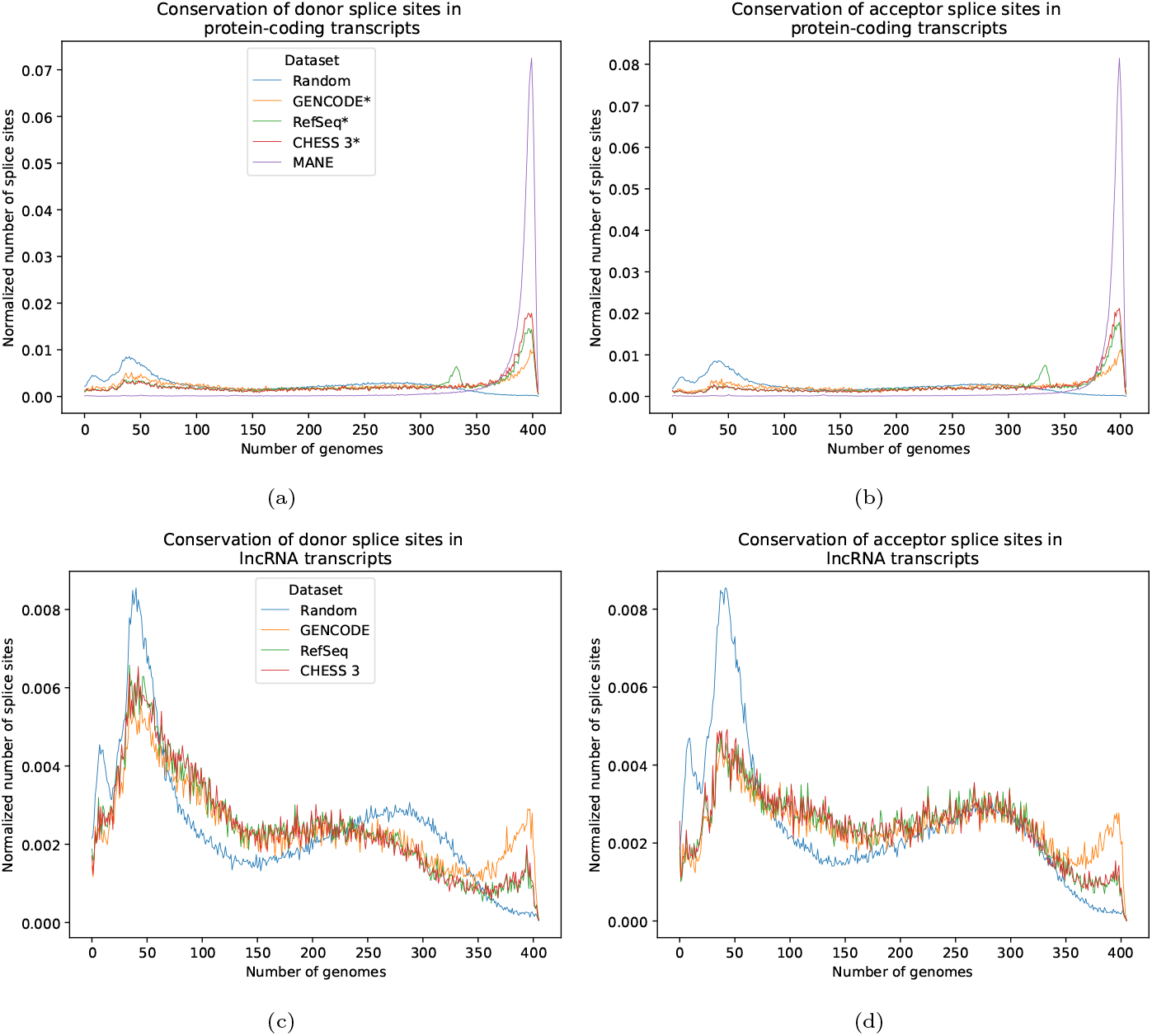
Distribution of the number of human splice sites with canonical dinucleotides (GT for donor and AG for acceptor sites) conserved in 405 mammals, computed for donor (a) and acceptor (b) sites of protein-coding genes, and donor (c) and acceptor (d) sites of lncRNAs. Each point shows a number of splice sites conserved (y-axis) in a given number of genomes (x-axis). Numbers are normalized by the total number of sites in the corresponding dataset in each category. The figure shows this statistic for annotations from GENCODE, RefSeq, CHESS 3, and MANE, as well as artificial splice sites (“Random”) generated from internal sequences of introns which are assumed to evolve neutrally. For protein-coding genes, we created subsets GENCODE, RefSeq, and CHESS 3, from which we removed MANE annotations because each of these datasets is a superset of MANE; the resulting datasets are designated as GENCODE^∗^, RefSeq^∗^, and CHESS 3^∗^ correspondingly.

We also calculated the most common species in which these splice sites are conserved, represented in Supplementary Table S2. These species mostly constitute primates, which suggests that their conservation is merely a result of having a relatively recent common ancestor with humans. These splice sites may be clade-specific, or they might represent erroneous annotations. We also calculated the same statistic for splice sites that have non-canonical dinucleotides on the introns’ flanks. Most of these splice sites constitute either U2-type sites flanked by GC-AG [57] or U12-type minor form introns [21, 22] flanked by the dinucleotides AT-AC. Supplementary Figure S1 shows these numbers; they follow the same pattern as splice sites with the canonical dinucleotides.

Given the striking pattern of conservation of the canonical dinucleotides of splice sites from MANE, we investigated the conservation of different positions around splice sites. Supplementary Figure S2 shows the pattern of conservation of bases as a function of their distance from the GT/AG splice site. As expected, the canonical dinucleotides (GT for donor sites and AG for acceptor sites) are the most conserved.

On the other hand, upstream positions for donor sites and downstream positions for acceptor sites show similar patterns of conservation. However, downstream positions for donor sites and upstream ones for acceptor sites are much less conserved, which is expected because these positions are intronic.

We further explored the question of how well splice sites are conserved at the human population level. Specifically, we calculated the fraction of splice sites having an SNP at a certain position, similar to the cross-species conservation of different positions shown in Supplementary Figure S2. To determine the presence of SNPs in the human population, we used the gnomAD database version 4.0.0 [8], focusing on loci that have at least one homozygous sample since a homozygous SNP at a splice site is very likely to cause incorrect splicing. Figure 3 shows these fractions, which we call “SNP rates,” calculated for each of the different gene catalogs. As expected, for protein-coding genes and their donor and acceptor splice sites, MANE has a much lower fraction of SNP sites at the canonical dinucleotides compared to random GT/AG positions, 0.2% versus 1.2%. On the other hand, splice sites in GENCODE^∗^ have only slightly lower SNP rates than randomly evolving sequences; RefSeq^∗^’s and CHESS 3^∗^’s rates are closer to MANE, but still somewhat higher. For lncRNAs from all of the catalogs, we observed that the SNP rates are relatively close to those of neutrally evolving sequences.

**Fig. 3.**
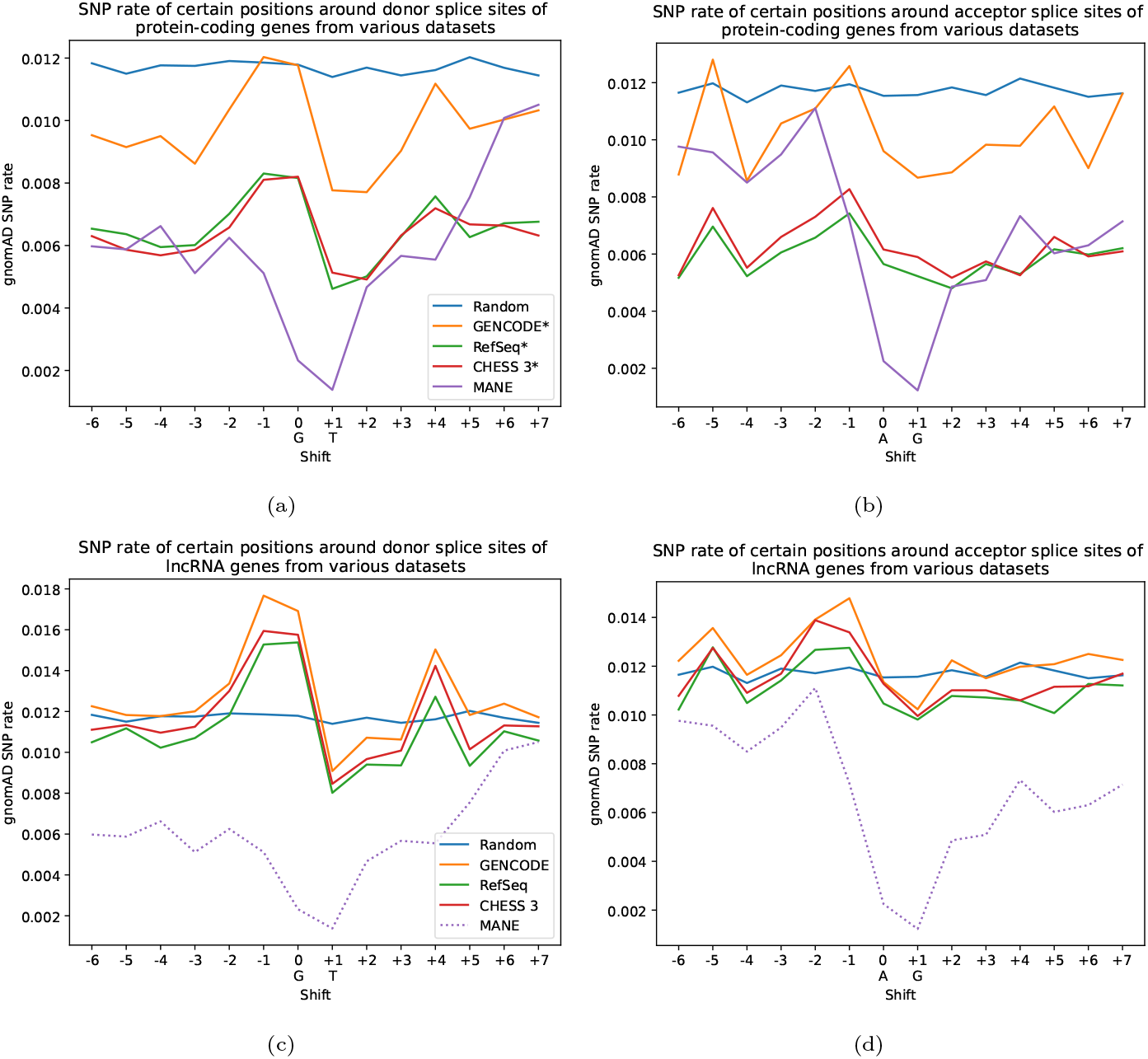
Rate of SNPs at positions near splice sites. Each point represents a proportion of splice sites from a certain dataset that have an SNP from the gnomAD dataset at a position either down- or upstream of the “canonical” dinucleotides. For example, for donor splice sites 0 is usually “G”, +1 is “T” (shown under the corresponding ticks on the horizontal axis), and -1 is the first nucleotide upstream of the splice site. We only considered SNPs that have one homozygous sample. Panels (a)-(b) show donor and acceptor sites of protein-coding genes, while (c)-(d) show values for donor and acceptor sites of lncRNAs. For lncRNAs, we included MANE sites from protein-coding genes as a baseline for splice sites under strong selection as MANE does not contain lncRNAs yet.

Our analysis suggests that some protein-coding splice sites and many more lncRNA splice sites include a subset of sites that are not under strong purifying selection. Otherwise, their average SNP rates should have been more similar to what we observed in splice sites from MANE. These sites might potentially represent non-functional and/or erroneous annotations.

### Classifying splice sites based on their conservation

Above we showed that splice sites from the major gene catalogs exhibit two clearly distinct patterns of conservation: MANE-like and random-like. For brevity, we refer to the former as “well-supported” and the latter as “less-supported.” We next decided to classify splice sites based on their conservation across species, and to compare their properties to see whether less-supported sites might be misannotated. To do so, we trained a binary classifier based on logistic regression that uses the number of species in which a certain position around a splice site is conserved; we trained models for donor and acceptor sites separately. We used the randomly generated sites as negative examples and the whole MANE database as positive ones, with 20% of the data set aside for testing; the Methods section contains a detailed description of the model.

Supplementary Figure S3 shows the receiver operating characteristic (ROC) curve illustrating the tradeoff between true positive and false positive rates for these models on the test data. We evaluated two types of models: one using only the conservation of the canonical dinucleotides GT/AG themselves, and the other one using the conservation of the other positions in the splicing motif (see the Methods section for more details). Both models show high accuracy on the test data with an area under the ROC curve (AUROC) measuring 0.974-0.979 for donor and acceptor sites. However, the full model has a slightly lower false positive rate for a given classification threshold, hence we chose it for further analyses. For classification, we used a threshold of 0.5 for the probability predicted by the regression model to classify sites as well-supported and less-supported. Given this threshold, the full models for donor and acceptor sites have F-scores of 0.933 and 0.949 correspondingly, see Supplementary Table S3.

In addition, we compared the probability output by the regression model to PhastCons [54] scores indicating whether a particular position in the genome is under negative selection. To do so, we used the PhastCons scores track available at the UCSC Genome Browser that were computed using the same 470-species alignment. For each site, we took a minimum of two PhastCons scores of the positions of their canonical dinucleotides (usually GT/AG) and computed the Pearson correlation between this quantity and the probability predicted by the regression model; this resulted in the Pearson correlation value of 0.8 across all sites from the datasets under consideration for which PhastCons scores were available.

We then applied the model to each dataset under consideration to label sites as well-supported or less-supported. Table 2 (columns 4 and 5) contains the number of donor and acceptor splice sites in each of the annotation databases classified by the model as well-supported. For protein-coding genes, we observed that in GENCODE^∗^ only 30% of donor and 40% of acceptor splices sites were well-supported according to the model, while for RefSeq^∗^ and CHESS 3^∗^, the proportion was higher, at 44% for donor sites and 46-49% for acceptor sites, suggesting the RefSeq and CHESS 3 has somewhat more reliable annotations of protein-coding transcripts. For lncRNAs, no more than 17% splice sites were classified as well-supported across all datasets. We observed similar results for non-canonical splice sites, these numbers are presented in Supplementary Table S4. Supplementary Figure S4 shows the relationship between the probability of a donor (acceptor) splice sites being classified as “well-supported” and the number of genomes in which the canonical dinucleotides of the particular splice-site are conserved in; most sites that have their dinucleotides conserved in less that 200 species are classified as “less-supported.”

Columns 6 and 7 of Table 2 also show the total number of transcripts in each annotation and the number of “well-supported transcripts” where each splice site is either shared with a transcript from MANE, or classified as “well-supported” by our model; we only considered transcripts with at least one intron present. We observed that for protein-coding genes,

GENCODE^∗^, RefSeq^∗^ and CHESS 3^∗^ 49%, 40% and 48% of transcripts have fully well-supported splice sites The number of well-supported lncRNAs is much lower in each dataset: 1-1.1% for RefSeq and CHESS 3 and 2% for GENCODE. Supplementary Figure S5 shows the number of well-supported transcripts in each gene type and dataset split by the number of introns in a transcript. We also broke down the number of well-supported and less-supported splice sites of protein-coding genes by whether they are located inside a MANE exon. As Supplementary Figure S2 shows, positions within the exon are conserved similarly to the canonical dinucleotides, and alternative splice sites located within exons could be mistakenly labeled as well-supported, resulting in more such sites. Supplementary Table S5 shows these numbers: there are nearly 10 times more splice sites outside of MANE exons overall, and the ones located inside exons are slightly more likely to be classified as “well-supported.” For all three datasets, around half of the splice sites inside MANE exons are well-supported. On the other hand, this statistic varies between different datasets for the sites outside of such exons. For example, CHESS 3^∗^ has 47-48% of such sites well-supported, RefSeq^∗^ has 42-46%, and GENCODE^∗^ has 25-27%.

We further compared SNP rates in the human population for well-supported and less-supported splice sites, again using the gnomAD human variation database and focusing on sites where at least one individual had a homozygous SNP. Figure 4 shows these rates for different datasets. For protein-coding genes (Figure 4, panels (a)-(b)), we observed that SNP rates for the canonical dinucleotides (positions 0 and +1) were 2–6 times lower for the well-supported subset (as predicted by our classifier) compared to its less-supported counterpart. We also note that the curves corresponding to less-supported sites are closer to the Random (neutrally evolving) sites, while well-supported sites in all three databases have SNP rates similar to MANE. However, for lncRNAs (Figure 4, panels (c)-(d)) the separation is a little less clear: although the less-supported sites have SNP rate pattern close to the Random ones, the well-supported sites have only 1.5-2 times smaller SNP rates at the canonical dinucleotides, and these rates are also much higher than the rates of protein-coding sites from MANE.

**Fig. 4.**
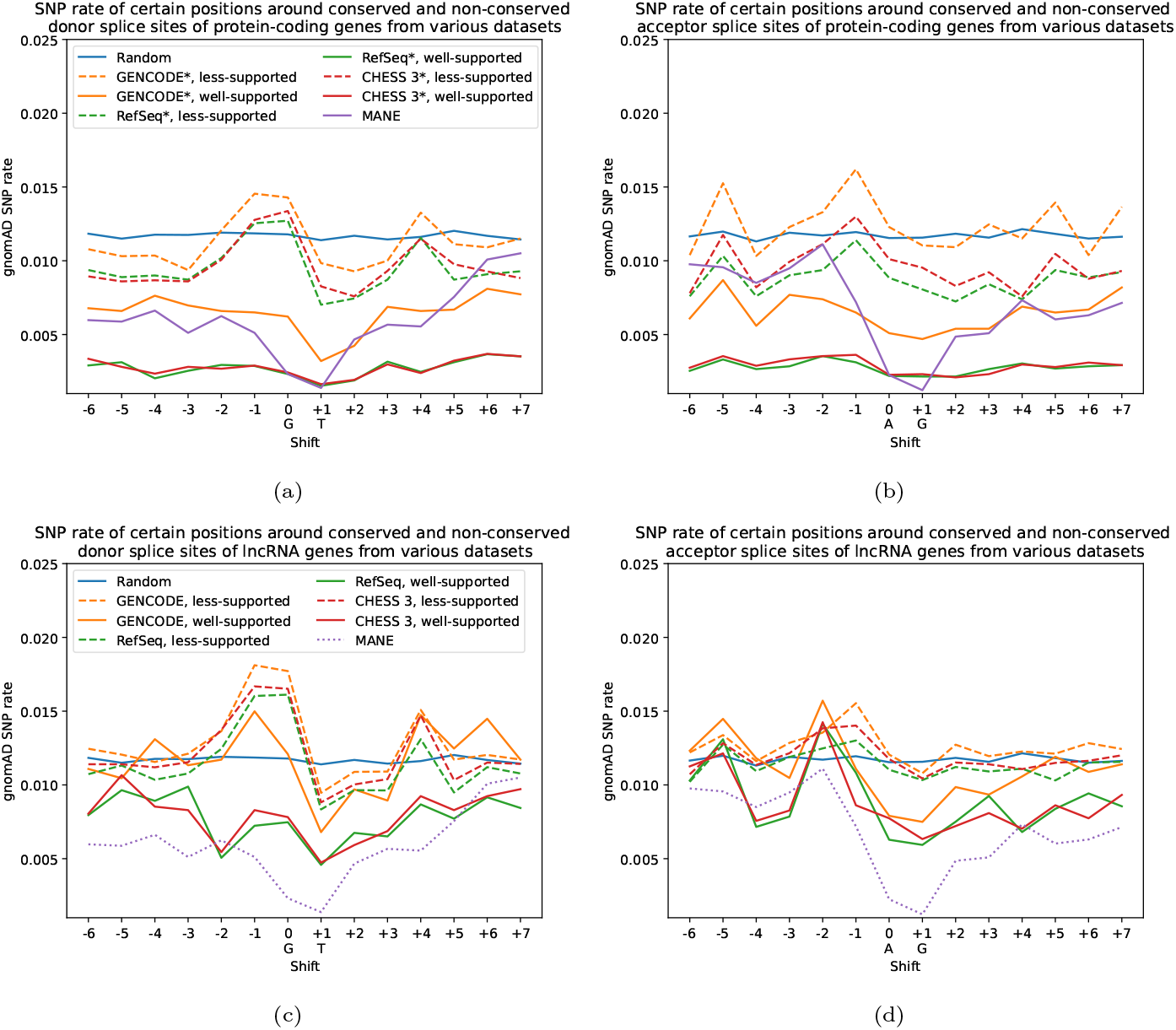
Rate of homozygous SNPs at positions near splice sites. Each point represents a proportion of splice sites from a certain dataset that have an SNP at a position either down- or upstream of the “canonical” dinucleotides. For example, for donor splice sites 0 is usually “G”, +1 is “T” (shown under the corresponding ticks on the horizontal axis), and -1 is the first nucleotide upstream of the splice site. We only considered SNPs from the gnomAD database that have at least one homozygous sample. Panels (a)-(b) show donor and acceptor sites of protein-coding genes, while (c)-(d) show values for donor and acceptor sites of lncRNAs. Solid lines represent subsets classified as “well-supported” by our model, while dashed ones correspond to “less-supported” splice sites. The dotted purple line on panels (c)-(d) represents the rate of SNPs for splice sites from protein-coding genes of MANE for comparison.

Apart from calculating the SNP rates, we compared the frequencies of homozygous SNPs overlapping the canonical dinucleotides of the splice sites classified by the model as either well-supported or less-supported. Supplementary Figure S6 shows these frequencies for different datasets. For donor and acceptor sites from protein-coding genes, the median frequencies of homozygous SNPs in well-supported sites are 2-3 times smaller than for less-supported ones. In addition, their interquartile range is 3-6 times smaller. This is also true for donor sites of lncRNAs from GENCODE, CHESS 3, as well as for acceptor sites of lncRNAs from GENCODE and RefSeq. Frequency distributions of well-supported and less-supported donor sites of lncRNAs from RefSeq and acceptor sites of lncRNAs from CHESS 3 are somewhat closer.

We also examined how many splice sites have SNPs overlapping their canonical dinucleotides associated with diseases. Table 3 shows the number and the fraction of splice sites relative to their total number in the respective dataset that have at least one SNP from the ClinVar database [31] classified as “pathogenic” or “likely pathogenic.” Well-supported splice sites of protein-coding genes have a two to three times higher ratio of potentially pathogenic SNPs than less-supported ones across all datasets. Well-supported sites of lncRNAs have a consistently low number of pathogenic SNPs, while less-supported ones from lncRNAs possess even fewer such variants. However, we note that non-coding regions are under-ascertained in clinical variant databases, as reported previously [13].

**Table 3.** Number of splice sites in each conservation category having an SNP classified as “pathogenic” and “likely pathogenic” from ClinVar database overlapping its canonical dinucleotides. Sites are classified as “well-supported” or “less-supported” by our model as per subsection 4.3. Numbers in parentheses show the percentage of the sites relative to the total number of sites in that category.

| Dataset | “Well-supported” sites |  | “Less-supported” sites |  |
| --- | --- | --- | --- | --- |
|  | Donor | Acceptor | Donor | Acceptor |
| *Protein-Coding* |  |  |  |  |
| GENCODE* | 87 (0.82%) | 80 (0.80%) | 53 (0.23%) | 36 (0.21%) |
| RefSeq* | 70 (0.25%) | 62 (0.24%) | 41 (0.12%) | 35 (0.12%) |
| CHESS 3* | 71 (0.30%) | 64 (0.28%) | 42 (0.16%) | 33 (0.15%) |
| *lncRNA* |  |  |  |  |
| GENCODE | 16 (0.20%) | 10 (0.10%) | 7 (0.01%) | 3 (0.01%) |
| RefSeq | 9 (0.22%) | 6 (0.10%) | 4 (0.01%) | 4 (0.01%) |
| CHESS 3 | 12 (0.28%) | 10 (0.18%) | 4 (0.01%) | 4 (0.01%) |

We also examined the connection between multiple species conservation and gene expression using RNA-seq data. Our model classifies each acceptor and donor splice sites as either well-supported or less-supported. This way, each intron consisting of a pair of a donor and an acceptor splice site belongs to either one of 4 categories: (1) neither site is well-supported (2) only the donor site is well-supported (3) only the acceptor site is well-supported (4) both sites are well-supported. We employed data from the GTEx project [10] that was assembled using StringTie2 [28] and was postprocessed by TieBrush [58] to obtain the junction coverage. This data was used to generate the CHESS 3 gene catalog, further technical details on the pipeline can be found in [59]. For each intron we calculated the number of reads supporting the particular donor-acceptor junction reflecting expression of isoforms using this intron. To integrate the data, we calculated the maximum coverage for each intron across all tissues; the breakdown for each individual tissue is available in Appendix, Supplementary Figures S7 and S8. Figure 5 shows distributions of read coverage between introns of different conservation categories, for both protein-coding in lncRNA genes. As the figure shows, introns with both sides that are well-supported have median max coverage that is 2-3 times higher than the introns that have at least one less-supported site, which can be observed for both protein-coding and lncRNA genes. Supplementary Figures S7 and S8 show coverage distribution across individual tissues that show the same pattern.

**Fig. 5.**
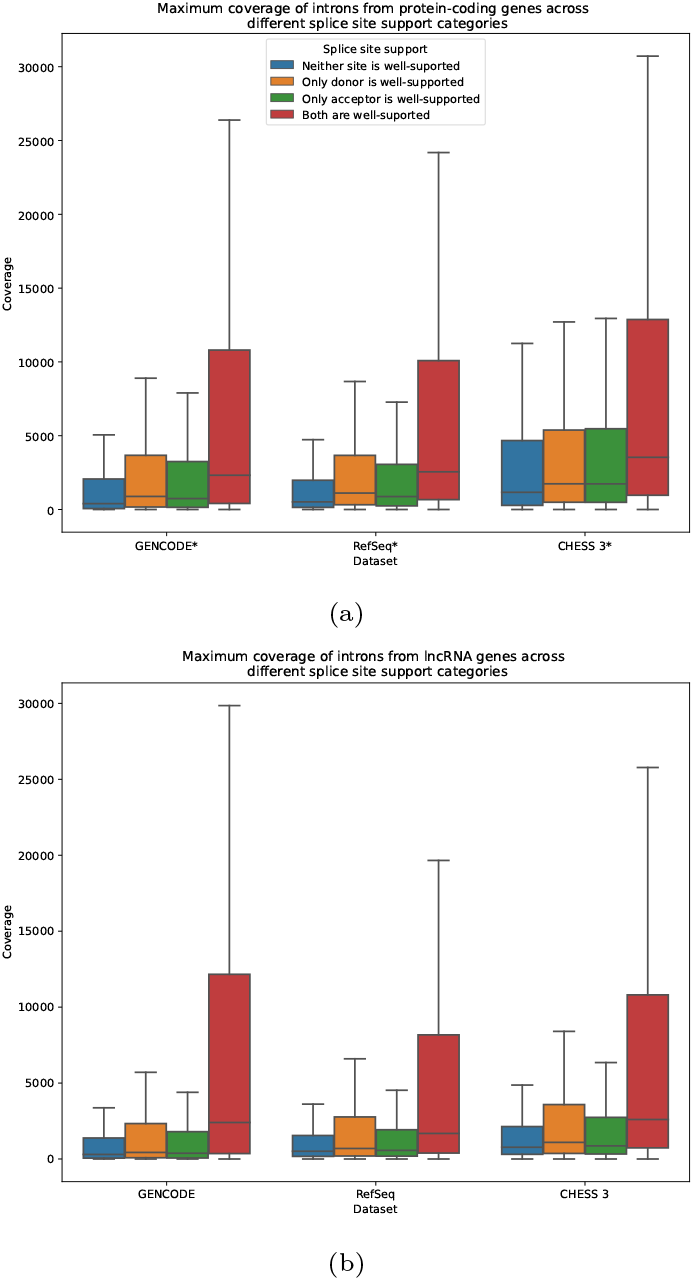
Box plots showing the maximum number of reads supporting exon junctions of certain types across all tissues from GTEx data. Panel (a) shows data for protein-coding genes (GENCODE^∗^, RefSeq^∗^, and CHESS 3^∗^ datasets), while panel (b) represents lncRNAs (GENCODE, RefSeq, CHESS 3). Each box plot shows the median (dashed red line), the interquartile range (solid top and bottom borders of the box), and minimum and maximum values within ±1.5 of the interquartile range (whiskers), outliers are not shown.

In addition, we explored how many isoforms of the same gene use well-supported and less-supported splice sites. In other words, for each splice site we computed the number of isoforms that use that particular site. Supplementary Figure S9 shows the distribution of these values in each gene catalog, for both donor and acceptor sites in protein-coding genes and lncRNAs. For protein-coding genes, well-supported splice sites are more likely to be shared between multiple isoforms. However, we did not observe a similar pattern in lncRNAs, except for donor sites from the GENCODE annotation that also showed a notable difference between well-supported and less-supported sites. We note that lncRNA genes have fewer isoforms overall, which might explain some of the disparity between protein-coding genes and lncRNAs.

### Case study: spotting potentially suspicious isoforms

To demonstrate the utility of our model, we manually inspected several isoforms that contain less-supported splice sites to see if they appear to be non-functional. In the following, we provide two examples, one for protein-coding genes and one for lncRNAs.

In particular, we looked at the heat shock protein family B (small) member 1, or the HSPB1 gene. Its MANE transcript with GENCODE ID ENST00000248553.7 contains 2 exons and produces a protein that is 205 amino acids long. One of the alternative transcripts with the ID ENST00000674547.1 from the GENCODE database differs from the MANE isoform by one donor site, which was marked as “less-supported” by our model. This alternative site results in a premature stop codon in the protein sequence, which yields a protein that is only 143 amino acids long, which is nearly 30% times shorter than its MANE isoform; Supplementary Figure S10a shows these two isoforms. In addition, the transcriptomic support for the splice junction containing the alternative site is also poor: tens of reads as opposed to millions or hundreds of thousands for the MANE isoform, depending on the tissue see Supplementary Table S6, columns 2 and 3, for the exact numbers. Given these data, we hypothesize that ENST00000674547.1 is either a technical artifact or a result of spurious transcription.

We also looked at CHASERR, a highly-conserved lncRNA located near the Chromodomain helicase DNA binding protein 2, a protein associated with a neurological disease [30]. Recently, CHASERR itself was shown to be critical for viability [48], and its deletion was associated with a neurological disorder [17]. One of its isoforms in GENCODE, with ID ENST00000557682.6, contains four introns, and all splice sites of those introns are “well-supported” according to our model. Transcripts with the same intron structure are also present in the RefSeq annotation. However, a GENCODE isoform with ID ENST00000653163.1 differs from the above-mentioned transcript by the location of the first donor site, which is located much further upstream: at position 92,819,999 of chromosome 15, as opposed to 92,883,187. The alternative donor site is “less-supported” according to our model and has only 2 reads covering that junction in the GTEx dataset. On the other hand, the intron junction of ENST00000557682.6 sharing the same acceptor site that has a well-supported donor site, is covered by thousands of reads across multiple tissues, see Supplementary Table S6, columns 4 and 5. Given that the isoform with a “less-supported” donor site occupies a locus 4.7 times longer and the longer intron has poor transcriptomic support, we believe that the longer transcript could be non-functional. Supplementary Figure S10b illustrates the structure of these two isoforms.

## Discussion

In this study, we found that the canonical dinucleotides from both donor and acceptor splice sites of the consensus MANE dataset exhibit a striking pattern of conservation: nearly all of them are conserved in more than 350 mammalian species. In contrast, splice sites from the leading gene catalogs – GENCODE, RefSeq, and CHESS 3 – that are not shared with MANE exhibit two different patterns of conservation. The first pattern resembles MANE, where the splice sites are conserved in more than 350 species, while the second one resembles neutrally evolving sequences, at both the micro- and macroevolutionary levels. To compare the properties of these two groups of splice sites, we trained a logistic regression model using the MANE dataset as the source of positive examples and using randomly chosen dinucleotide sites from within introns to represent (albeit imperfectly) neutrally evolving sequences. We then applied this model to the rest of the GENCODE, RefSeq, and CHESS gene catalogs excluding MANE to classify splice sites as either well-supported or less-supported.

We found that 30-50% of splice sites from coding genes and less than 17% of splice sites from lncRNA can be classified as well-supported. Splice sites classified as less-supported had SNP rates in the human population that were consistent with neutrally evolving sequences, while well-supported ones had patterns of SNP rates and frequencies similar to MANE. In addition, we observed that introns where both splice sites are well-supported have better transcriptomic support. We also found that less-supported splice sites are less likely to be shared by different isoforms of the same gene. We calculated the number of transcripts whose splice sites were either classified as well-supported by our model or shared with a transcript from MANE. For protein-coding genes, 41%-56% belong to this category, and only 0.5-2% have all their sites well-supported. These transcripts can be used as a high-confidence subset of the gene catalogs we studied.

Our findings are consistent with the previous studies of splice site evolution. For example, it was observed before that genes that are highly conserved have higher expression levels [20, 6] and conserved exons are more likely to be included in multiple transcripts [35]. Other studies [29, 53] found that splice sites that are not conserved in other species are more likely to carry disruptive SNPs in their motifs, and transcripts enriched in such variants have lower expression levels [11]; it was also observed that splice sites of lncRNAs are less conserved than ones of mRNAs [60].

Some previous studies also found a lack of conservation of some splice sites using whole genome alignments [51, 52], but they assumed that these patterns arose due to alignment errors. Using the large gnomAD collection of human variation, we were able to examine SNP rates in the human population and show that splice sites that are less-supported also have higher SNP rates and frequencies. This finding suggests that the majority of splice sites lacking conservation across species are simply not under selection in the human population as opposed to being poorly aligned.

This study has several limitations. First, unlike some previous studies of the evolutionary dynamics of alternative splicing [3, 33, 16], we rely purely on the conservation of DNA sequences without taking into account whether a conserved splice site in a non-human genome is actually functional (information that is usually not known). Unfortunately, the incomplete status of many other genome annotations prevented us from incorporating them into our analysis. Second, we realize that the training data we used for our model could introduce biases. For example, the MANE dataset was constructed by choosing one “best” isoform per protein-coding gene, and conservation was one of the criteria. This could potentially contribute to the stronger conservation signal we observed in that data. In addition, randomly chosen dinucleotide sequences from the interior of introns might not be the ideal choice for neutrally evolving sequences. We also note that MANE might not be an appropriate baseline for comparison for non-coding genes. However, the subset of experimentally verified lncRNAs is small, which makes it challenging to create a training set based on these genes. At the same time, in our study at least a subset of lncRNAs showed levels of conservation (at their splice sites) comparable to protein-coding genes. We believe that our model can be still useful for such genes; e.g., if most splice sites of a lncRNA transcript are well-supported, then less-supported ones should be dealt with caution.

We hope this study will help improve human genome annotation by demonstrating the utility of using large-scale evolutionary conservation for functional annotation of splicing. According to our analysis, highly-conserved splice sites from MANE alone constitute at least 75% of all the splice sites in protein-coding isoforms in all genome annotations, and together with their well-supported counterparts from the complementary subsets account for 80-90% of all splice sites (Table 2). This finding is in concordance with previous studies showing that MANE and APPRIS transcripts represent the most biologically and clinically relevant isoforms [42, 43].

Hence, we believe that splice site conservation should be an important factor in constructing a genome annotation. However, only a few methods currently use this information directly, either for annotation or splice site prediction [49]. A common data structure used in RNA assembly called splice graph was generalized to integrate sequence homology information between species [61], but it was used for finding clusters of orthologous exons and yet to be employed for RNA-seq assembly. As higher-quality genomes along with their alignments become available, conservation-based methods have the potential to be a powerful aid in constructing functional annotations. However, despite the recent advances in the field of alignment [34, 2, 27], our analysis shows that even the most complete whole-genome alignments to date miss many alignments of human exons, and further progress in this area is needed to improve the completeness of the alignments.

We also highlighted a subset of splice sites and corresponding isoforms in the leading human annotation catalogs that appear to be under strong selection. This subset can be used as a high-confidence representation of the annotation. At the same time, less-supported splice sites might require further scrutiny since splicing could be inherently error-prone [24, 23, 40, 19, 4]. This hypothesis is backed up by the fact that well-supported splice sites have higher SNP rates and frequencies in the human population consistent with randomly selected sites, which suggests that they are not under as strong negative selection. In addition, a recent study using proteomics analysis found that only 1 in 6 alternative isoforms were predicted to be functional [41]. However, pinpointing exactly which splice sites are errors would require further study incorporating extra data. Here we have focused on the human genome because it has the highest-quality annotation, but in the future, we hope to extend our analysis to the annotations of other species.

## Data availability

The data and the code of the model are available at GitHub via URL: https://github.com/iminkin/splice-sites-conservation. This repository, including all the code and the resulting data, was archived at Zenodo and is available with the following DOI: 10.5281/zenodo.14893716.

## Acknowledgements

We would like to thank Mihaela Pertea, Aleksey Zimin, Ales Varabyou, Beril Erdogdu, and Kuan-Hao Chao for useful discussions and suggestions.

## Author contributions

I.M. and S.S. conceptualized the project. I.M. performed the investigation, formal analysis, and wrote the original draft. Both authors edited and revised all versions of the manuscript.

## References

1. Paulo Amaral, Silvia Carbonell-Sala, Francisco M De La Vega, Tiago Faial, Adam Frankish, Thomas Gingeras, Roderic Guigo, Jennifer L Harrow, Artemis G Hatzigeorgiou, Rory Johnson, et al. The status of the human gene catalogue. Nature, 622(7981):41–47, 2023.

2. Joel Armstrong, Glenn Hickey, Mark Diekhans, Ian T Fiddes, Adam M Novak, Alden Deran, Qi Fang, Duo Xie, Shaohong Feng, Josefin Stiller, et al. Progressive cactus is a multiple-genome aligner for the thousand-genome era. Nature, 587(7833):246–251, 2020.

3. Nuno L Barbosa-Morais, Manuel Irimia, Qun Pan, Hui Y Xiong, Serge Gueroussov, Leo J Lee, Valentina Slobodeniuc, Claudia Kutter, Stephen Watt, Recep Colak, et al. The evolutionary landscape of alternative splicing in vertebrate species. Science, 338(6114):1587–1593, 2012.

4. Florian Bénitières, Anamaria Necsulea, and Laurent Duret. Random genetic drift sets an upper limit on mrna splicing accuracy in metazoans. Elife, 13:RP93629, 2024.

5. Mathieu Blanchette, W James Kent, Cathy Riemer, Laura Elnitski, Arian FA Smit, Krishna M Roskin, Robert Baertsch, Kate Rosenbloom, Hiram Clawson, Eric D Green, et al. Aligning multiple genomic sequences with the threaded blockset aligner. Genome research, 14(4):708– 715, 2004.

6. Benjamin J Blencowe. Alternative splicing: new insights from global analyses. Cell, 126(1):37–47, 2006.

7. Massimo Cavallaro, Mark D Walsh, Matt Jones, James Teahan, Simone Tiberi, Bärbel Finkenstädt, and Daniel Hebenstreit. 3’-5’ crosstalk contributes to transcriptional bursting. Genome biology, 22:1–20, 2021.

8. Siwei Chen, Laurent C Francioli, Julia K Goodrich, Ryan L Collins, Masahiro Kanai, Qingbo Wang, Jessica Alföldi, Nicholas A Watts, Christopher Vittal, Laura D Gauthier, et al. A genomic mutational constraint map using variation in 76,156 human genomes. Nature, pages 1–11, 2023.

9. Peter JA Cock, Tiago Antao, Jeffrey T Chang, Brad A Chapman, Cymon J Cox, Andrew Dalke, Iddo Friedberg, Thomas Hamelryck, Frank Kauff, Bartek Wilczynski, et al. Biopython: freely available python tools for computational molecular biology and bioinformatics. Bioinformatics, 25(11):1422–1423, 2009.

10. GTEx Consortium, Kristin G Ardlie, David S Deluca, Ayellet V Segrè, Timothy J Sullivan, Taylor R Young, Ellen T Gelfand, Casandra A Trowbridge, Julian B Maller, Taru Tukiainen, et al. The genotype-tissue expression (gtex) pilot analysis: multitissue gene regulation in humans. Science, 348(6235):648–660, 2015.

11. Stepan V Denisov, Georgii A Bazykin, Roman Sutormin, Alexander V Favorov, Andrey A Mironov, Mikhail S Gelfand, and Alexey S Kondrashov. Weak negative and positive selection and the drift load at splice sites. Genome biology and evolution, 6(6):1437–1447, 2014.

12. W Ford Doolittle. Genes in pieces: were they ever together? Nature, 272(5654):581–582, 1978.

13. Jamie M Ellingford, Joo Wook Ahn, Richard D Bagnall, Diana Baralle, Stephanie Barton, Chris Campbell, Kate Downes, Sian Ellard, Celia Duff-Farrier, David R FitzPatrick, et al. Recommendations for clinical interpretation of variants found in non-coding regions of the genome. Genome medicine, 14(1):73, 2022.

14. Jessica H Fong, Terence D Murphy, and Kim D Pruitt. Comparison of refseq protein-coding regions in human and vertebrate genomes. BMC genomics, 14:1–16, 2013.

15. Adam Frankish, Mark Diekhans, Irwin Jungreis, Julien Lagarde, Jane E Loveland, Jonathan M Mudge, Cristina Sisu, James C Wright, Joel Armstrong, If Barnes, et al. Gencode 2021. Nucleic acids research, 49(D1):D916–D923, 2021.

16. Andreas Franz, A Ioana Weber, Marco Preußner, Nicole Dimos, Alexander Stumpf, Yanlong Ji, Laura Moreno-Velasquez, Anne Voigt, Frederic Schulz, Alexander Neumann, et al. Branch point strength controls species-specific camk2b alternative splicing and regulates ltp. Life science alliance, 6(3), 2023.

17. Vijay S. Ganesh, Kevin Riquin, Nicolas Chatron, Esther Yoon, Kay-Marie Lamar, Miriam C. Aziz, Pauline Monin, Melanie C. O’Leary, Julia K. Goodrich, Kiran V. Garimella, Eleina England, Ben Weisburd, François Aguet, Carlos A. Bacino, David R. Murdock, Hongzheng Dai, Jill A. Rosenfeld, Lisa T. Emrick, Shamika Ketkar, Yael Sarusi, Damien Sanlaville, Saima Kayani, Brian Broadbent, Alisée Pengam, Bertrand Isidor, Stéphane Bezieau, Benjamin Cogné, Daniel G. MacArthur, Igor Ulitsky, Gemma L. Carvill, and Anne O’Donnell-Luria. Neurodevelopmental disorder caused by deletion of ¡i¿chaserr¡/i¿, a lncrna gene. New England Journal of Medicine, 391(16):1511–1518, 2024.

18. W Gilbert. Why genes in pieces? Nature, 271(5645):501, 1978.

19. Jean-François Gout, W Kelley Thomas, Zachary Smith, Kazufusa Okamoto, and Michael Lynch. Large-scale detection of in vivo transcription errors. Proceedings of the National Academy of Sciences, 110(46):18584–18589, 2013.

20. Philip Green, David Lipman, LaDeana Hillier, Robert Waterston, David States, and Jean-Michel Claverie. Ancient conserved regions in new gene sequences and the protein databases. Science, 259(5102):1711–1716, 1993.

21. Stephen L. Hall and Richard A. Padgett. Conserved sequences in a class of rare eukaryotic nuclear introns with non-consensus splice sites. Journal of Molecular Biology, 239(3):357–365, 1994.

22. Stephen L. Hall and Richard A. Padgett. Requirement of u12 snrna for in vivo splicing of a minor class of eukaryotic nuclear pre-mrna introns. Science, 271(5256):1716–1718, 1996.

23. Shu-Ning Hsu and Klemens J Hertel. Spliceosomes walk the line: splicing errors and their impact on cellular function. RNA biology, 6(5):526–530, 2009.

24. Laurence D Hurst. Evolutionary genomics and the reach of selection. Journal of Biology, 8:1–5, 2009.

25. Manuel Irimia and Scott William Roy. Origin of spliceosomal introns and alternative splicing. Cold Spring Harbor perspectives in biology, 6(6):a016071, 2014.

26. W James Kent, Charles W Sugnet, Terrence S Furey, Krishna M Roskin, Tom H Pringle, Alan M Zahler, and David Haussler. The human genome browser at ucsc. Genome research, 12(6):996–1006, 2002.

27. Bryce Kille, Advait Balaji, Fritz J Sedlazeck, Michael Nute, and Todd J Treangen. Multiple genome alignment in the telomere-to-telomere assembly era. Genome Biology, 23(1):182, 2022.

28. Sam Kovaka, Aleksey V Zimin, Geo M Pertea, Roham Razaghi, Steven L Salzberg, and Mihaela Pertea. Transcriptome assembly from long-read rna-seq alignments with stringtie2. Genome biology, 20:1–13, 2019.

29. Yerbol Z Kurmangaliyev, Roman A Sutormin, Sergey A Naumenko, Georgii A Bazykin, and Mikhail S Gelfand. Functional implications of splicing polymorphisms in the human genome. Human Molecular Genetics, 22(17):3449– 3459, 2013.

30. Kay-Marie J Lamar and Gemma L Carvill. Chromatin remodeling proteins in epilepsy: lessons from chd2-associated epilepsy. Frontiers in molecular neuroscience, 11:208, 2018.

31. Melissa J Landrum, Jennifer M Lee, George R Riley, Wonhee Jang, Wendy S Rubinstein, Deanna M Church, and Donna R Maglott. Clinvar: public archive of relationships among sequence variation and human phenotype. Nucleic acids research, 42(D1):D980–D985, 2014.

32. Michael Lynch and Aaron O Richardson. The evolution of spliceosomal introns. Current opinion in genetics & development, 12(6):701–710, 2002.

33. Jason Merkin, Caitlin Russell, Ping Chen, and Christopher B Burge. Evolutionary dynamics of gene and isoform regulation in mammalian tissues. Science, 338(6114):1593–1599, 2012.

34. Ilia Minkin and Paul Medvedev. Scalable multiple whole-genome alignment and locally collinear block construction with sibeliaz. Nature communications, 11(1):6327, 2020.

35. Barmak Modrek and Christopher J Lee. Alternative splicing in the human, mouse and rat genomes is associated with an increased frequency of exon creation and/or loss. Nature genetics, 34(2):177–180, 2003.

36. Joannella Morales, Shashikant Pujar, Jane E Loveland, Alex Astashyn, Ruth Bennett, Andrew Berry, Eric Cox, Claire Davidson, Olga Ermolaeva, Catherine M Farrell, et al. A joint ncbi and embl-ebi transcript set for clinical genomics and research. Nature, 604(7905):310–315, 2022.

37. Nuala A O’Leary, Mathew W Wright, J Rodney Brister, Stacy Ciufo, Diana Haddad, Rich McVeigh, Bhanu Rajput, Barbara Robbertse, Brian Smith-White, Danso Ako-Adjei, et al. Reference sequence (refseq) database at ncbi: current status, taxonomic expansion, and functional annotation. Nucleic acids research, 44(D1):D733–D745, 2016.

38. F. Pedregosa, G. Varoquaux, A. Gramfort, V. Michel, B. Thirion, O. Grisel, M. Blondel, P. Prettenhofer, R. Weiss, V. Dubourg, J. Vanderplas, A. Passos, D. Cournapeau, M. Brucher, M. Perrot, and E. Duchesnay. Scikit-learn: Machine learning in Python. Journal of Machine Learning Research, 12:2825–2830, 2011.

39. Mihaela Pertea, Alaina Shumate, Geo Pertea, Ales Varabyou, Florian P Breitwieser, Yu-Chi Chang, Anil K Madugundu, Akhilesh Pandey, and Steven L Salzberg. Chess: a new human gene catalog curated from thousands of large-scale rna sequencing experiments reveals extensive transcriptional noise. Genome biology, 19(1):1–14, 2018.

40. Joseph K Pickrell, Athma A Pai, Yoav Gilad, and Jonathan K Pritchard. Noisy splicing drives mrna isoform diversity in human cells. PLoS genetics, 6(12):e1001236, 2010.

41. Fernando Pozo, Laura Martinez-Gomez, Thomas A Walsh, José Rodriguez, Tomas Di Domenico, Federico Abascal, Jesús Vazquez, and Michael L Tress. Assessing the functional relevance of splice isoforms. NAR Genomics and Bioinformatics, 3(2):qab044, 2021.

42. Fernando Pozo, José Manuel Rodriguez, Laura Martìnez Gómez, Jesus Vazquez, and Michael L Tress. Appris principal isoforms and mane select transcripts define reference splice variants. Bioinformatics, 38(Supplement 2):ii89–ii94, 2022.

43. Fernando Pozo, José Manuel Rodriguez, Jesús Vazquez, and Michael L Tress. Clinical variant interpretation and biologically relevant reference transcripts. NPJ Genomic Medicine, 7(1):59, 2022.

44. Arjun Raj, Charles S Peskin, Daniel Tranchina, Diana Y Vargas, and Sanjay Tyagi. Stochastic mrna synthesis in mammalian cells. PLoS biology, 4(10):e309, 2006.

45. Brian J Raney, Galt P Barber, Anna Benet-Pagès, Jonathan Casper, Hiram Clawson, Melissa S Cline, Mark Diekhans, Clayton Fischer, Jairo Navarro Gonzalez, Glenn Hickey, et al. The ucsc genome browser database: 2024 update. Nucleic Acids Research, 52(D1):D1082–D1088, 2024.

46. Jose Manuel Rodriguez, Fernando Pozo, Daniel Cerdán-Vélez, Tomás Di Domenico, Jesús Vázquez, and Michael L Tress. Appris: selecting functionally important isoforms. Nucleic Acids Research, 50(D1):D54–D59, 11 2021.

47. Igor B Rogozin, Liran Carmel, Miklos Csuros, and Eugene V Koonin. Origin and evolution of spliceosomal introns. Biology direct, 7:1–28, 2012.

48. Aviv Rom, Liliya Melamed, Noa Gil, Micah Jonathan Goldrich, Rotem Kadir, Matan Golan, Inbal Biton, Rotem Ben-Tov Perry, and Igor Ulitsky. Regulation of chd2 expression by the chaserr long noncoding rna gene is essential for viability. Nature Communications, 10(1):5092, 2019.

49. Dominic Rose, Michael Hiller, Katharina Schutt, Jörg Hackermüller, Rolf Backofen, and Peter F Stadler. Computational discovery of human coding and non-coding transcripts with conserved splice sites. Bioinformatics, 27(14):1894–1900, 2011.

50. Valerie A Schneider, Tina Graves-Lindsay, Kerstin Howe, Nathan Bouk, Hsiu-Chuan Chen, Paul A Kitts, Terence D Murphy, Kim D Pruitt, Françoise Thibaud-Nissen, Derek Albracht, et al. Evaluation of grch38 and de novo haploid genome assemblies demonstrates the enduring quality of the reference assembly. Genome research, 27(5):849–864, 2017.

51. Virag Sharma, Anas Elghafari, and Michael Hiller. Coding exon-structure aware realigner (CESAR) utilizes genome alignments for accurate comparative gene annotation. Nucleic Acids Research, 44(11):e103–e103, 03 2016.

52. Virag Sharma, Peter Schwede, and Michael Hiller. CESAR 2.0 substantially improves speed and accuracy of comparative gene annotation. Bioinformatics, 33(24):3985–3987, 08 2017.

53. Makoto K Shimada, Yosuke Hayakawa, Jun-ichi Takeda, Takashi Gojobori, and Tadashi Imanishi. A comprehensive survey of human polymorphisms at conserved splice dinucleotides and its evolutionary relationship with alternative splicing. BMC Evolutionary Biology, 10(1):1– 12, 2010.

54. Adam Siepel, Gill Bejerano, Jakob S Pedersen, Angie S Hinrichs, Minmei Hou, Kate Rosenbloom, Hiram Clawson, John Spieth, LaDeana W Hillier, Stephen Richards, et al. Evolutionarily conserved elements in vertebrate, insect, worm, and yeast genomes. Genome research, 15(8):1034– 1050, 2005.

55. Martin Šošić and Mile Šikić. Edlib: a c/c++ library for fast, exact sequence alignment using edit distance. Bioinformatics, 33(9):1394–1395, 2017.

56. Kevin Struhl. Transcriptional noise and the fidelity of initiation by rna polymerase ii. Nature structural & molecular biology, 14(2):103–105, 2007.

57. T. A. Thanaraj and Francis Clark. Human gc-ag alternative intron isoforms with weak donor sites show enhanced consensus at acceptor exon positions. Nucleic Acids Research, 29(12):2581–2593, 06 2001.

58. Ales Varabyou, Geo Pertea, Christopher Pockrandt, and Mihaela Pertea. Tiebrush: an efficient method for aggregating and summarizing mapped reads across large datasets. Bioinformatics, 37(20):3650–3651, 2021.

59. Ales Varabyou, Markus J Sommer, Beril Erdogdu, Ida Shinder, Ilia Minkin, Kuan-Hao Chao, Sukhwan Park, Jakob Heinz, Christopher Pockrandt, Alaina Shumate, et al. Chess 3: an improved, comprehensive catalog of human genes and transcripts based on large-scale expression data, phylogenetic analysis, and protein structure. Genome biology, 24(1):249, 2023.

60. Stefan Washietl, Manolis Kellis, and Manuel Garber. Evolutionary dynamics and tissue specificity of human long noncoding rnas in six mammals. Genome research, 24(4):616–628, 2014.

61. Diego Javier Zea, Sofya Laskina, Alexis Baudin, Hugues Richard, and Elodie Laine. Assessing conservation of alternative splicing with evolutionary splicing graphs. Genome Research, 31(8):1462–1473, 2021.

