## Supplementary Information for "Conservation assessment of human splice site annotation based on a 470-genome alignment"

**Ilia Minkin**
Department of Biomedical Engineering
Center for Computational Biology
Johns Hopkins University, Baltimore, MD 21211, USA


**Steven L. Salzberg**
Department of Biomedical Engineering
Center for Computational Biology
Department of Computer Science
Department of Biostatistics
Johns Hopkins University, Baltimore, MD 21211, USA


| Chromosome | Aligned exons/genomes | Missing exons/genomes | Recovered exons/genomes |
|---|---|---|---|
| 1 | 55,324,093 | 8,287,637 | 4,749,605 |
| 2 | 42,977,856 | 5,285,994 | 3,294,472 |
| 3 | 30,816,387 | 3,108,843 | 2,003,824 |

Supplementary Table S1: The number of aligned exons from the generated "Random" dataset in the original alignment of 470 mammals restricted to 405 species (column 2, defined as $s_a$ in the main text), missing in the alignment (column 3, defined as $s_m$ in the main text), and recovered using our synteny-based realignment procedure (column 4, defined $s_r$ in the main text).

| | Donor sites | | | | Acceptor sites | | | |
|---|---|---|---|---|---|---|---|---|
| Organism | GENCODE$^{(*)}$ | RefSeq$^{(*)}$ | CHESS 3$^{(*)}$ | Random | GENCODE$^{(*)}$ | RefSeq$^{(*)}$ | CHESS 3$^{(*)}$ | Random |
| *Protein-Coding* | | | | | | | | |
| Chimpanzee | 88.07% | 88.50% | 87.92% | 85.68% | 87.61% | 88.03% | 87.21% | 86.00% |
| Pygmy chimpanzee | 87.80% | 88.09% | 87.47% | 85.69% | 87.15% | 87.47% | 86.61% | 85.96% |
| Western lowland gorilla | 84.44% | 84.93% | 83.91% | 82.96% | 84.02% | 84.55% | 82.96% | 83.16% |
| Sumatran orangutan | 80.09% | 80.87% | 80.34% | 79.30% | 81.47% | 81.33% | 80.32% | 79.41% |
| Northern white-cheeked gibbon | 76.75% | 76.89% | 76.13% | 75.79% | 78.32% | 77.82% | 76.91% | 75.79% |
| Silvery gibbon | 76.34% | 76.16% | 75.24% | 75.48% | 77.95% | 77.11% | 76.42% | 75.55% |
| Crab-eating macaque | 68.45% | 69.37% | 68.90% | 68.57% | 71.33% | 72.44% | 71.94% | 68.57% |
| Rhesus monkey | 68.52% | 69.57% | 69.07% | 68.50% | 71.36% | 72.42% | 71.91% | 68.40% |
| Gelada | 68.63% | 69.22% | 68.89% | 68.53% | 71.40% | 71.79% | 71.45% | 68.47% |
| Japanese macaque | 68.12% | 68.76% | 68.24% | 67.97% | 70.81% | 71.57% | 71.00% | 67.99% |
| Golden snub-nosed monkey | 68.13% | 69.16% | 68.99% | 67.83% | 70.69% | 71.60% | 71.24% | 67.71% |
| Green monkey | 68.34% | 67.56% | 68.46% | 67.71% | 70.41% | 69.47% | 70.54% | 67.71% |
| Olive baboon | 67.48% | 67.62% | 66.78% | 67.56% | 70.01% | 70.15% | 69.52% | 67.58% |
| Mandrill | 67.47% | 67.95% | 67.54% | 67.14% | 69.90% | 70.30% | 69.62% | 67.12% |
| Ugandan red Colobus | 67.24% | 67.85% | 67.29% | 67.18% | 69.52% | 69.95% | 69.33% | 67.10% |
| Red shanked douc langur | 67.34% | 67.90% | 67.54% | 67.09% | 69.78% | 70.07% | 69.64% | 66.89% |
| Red guenon | 67.31% | 67.67% | 67.16% | 67.00% | 69.67% | 69.68% | 68.90% | 67.15% |
| Allen's swamp monkey | 67.33% | 67.65% | 67.13% | 66.91% | 69.58% | 69.89% | 69.22% | 66.86% |
| Francois's langur | 67.36% | 67.81% | 67.31% | 66.46% | 69.96% | 70.28% | 69.94% | 66.32% |
| Mona monkey | 67.12% | 67.33% | 66.85% | 66.68% | 69.25% | 69.32% | 68.84% | 66.64% |
| De Brazza's monkey | 66.51% | 66.76% | 66.49% | 66.35% | 68.53% | 69.12% | 68.41% | 66.35% |
| Hanuman langur | 66.30% | 66.81% | 66.17% | 66.05% | 68.32% | 68.57% | 67.84% | 65.98% |
| Proboscis monkey | 50.59% | 50.23% | 50.26% | 54.78% | 53.07% | 52.74% | 52.23% | 54.70% |
| White-faced saki | 52.79% | 52.75% | 52.50% | 52.67% | 55.39% | 56.22% | 55.44% | 52.72% |
| Black-handed spider monkey | 52.55% | 52.22% | 52.49% | 52.74% | 54.82% | 55.62% | 54.86% | 52.90% |
| Mantled howler monkey | 52.17% | 52.23% | 52.16% | 52.21% | 54.16% | 55.53% | 54.53% | 52.14% |
| Tufted capuchin | 51.53% | 51.47% | 51.16% | 52.17% | 54.68% | 54.91% | 54.14% | 52.33% |
| White-tufted-ear marmoset | 51.41% | 51.17% | 50.84% | 51.62% | 54.22% | 54.44% | 53.80% | 51.70% |
| Bolivian titi | 51.11% | 50.91% | 50.70% | 51.68% | 54.17% | 54.48% | 53.59% | 51.82% |
| White-fronted capuchin | 50.56% | 50.45% | 50.00% | 51.56% | 53.51% | 54.12% | 53.03% | 51.71% |
| *lncRNA* | | | | | | | | |
| Chimpanzee | 90.07% | 89.73% | 89.89% | - | 89.73% | 89.98% | 89.59% | - |
| Pygmy chimpanzee | 89.70% | 89.51% | 89.52% | - | 89.48% | 90.04% | 89.43% | - |
| Western lowland gorilla | 87.69% | 87.14% | 87.37% | - | 87.60% | 88.11% | 87.45% | - |
| Sumatran orangutan | 81.28% | 81.50% | 81.44% | - | 83.30% | 83.93% | 83.62% | - |
| Northern white-cheeked gibbon | 75.58% | 75.80% | 75.61% | - | 77.34% | 77.82% | 77.28% | - |
| Silvery gibbon | 74.88% | 75.21% | 74.91% | - | 76.33% | 77.18% | 76.55% | - |
| Rhesus monkey | 68.41% | 68.21% | 68.11% | - | 71.37% | 71.80% | 71.10% | - |
| Crab-eating macaque | 68.46% | 68.24% | 68.00% | - | 71.34% | 71.80% | 71.05% | - |
| Gelada | 68.23% | 67.93% | 67.87% | - | 71.10% | 71.65% | 70.95% | - |
| Japanese macaque | 68.00% | 67.63% | 67.66% | - | 70.80% | 71.14% | 70.53% | - |
| Golden snub-nosed monkey | 67.92% | 67.63% | 67.60% | - | 70.60% | 71.14% | 70.46% | - |
| Green monkey | 67.74% | 66.42% | 67.45% | - | 70.77% | 70.12% | 70.37% | - |
| Red shanked douc langur | 67.37% | 66.94% | 66.98% | - | 70.12% | 70.35% | 69.97% | - |
| Olive baboon | 67.36% | 66.69% | 66.83% | - | 70.33% | 70.40% | 69.73% | - |
| Mandrill | 67.12% | 66.55% | 66.62% | - | 69.72% | 70.47% | 69.77% | - |
| Red guenon | 66.80% | 66.69% | 66.68% | - | 69.79% | 70.30% | 69.37% | - |
| Francois's langur | 66.88% | 66.38% | 66.50% | - | 69.69% | 69.56% | 69.13% | - |
| Allen's swamp monkey | 66.43% | 66.31% | 66.10% | - | 69.13% | 69.75% | 69.01% | - |
| Ugandan red Colobus | 66.50% | 66.07% | 66.02% | - | 69.17% | 69.43% | 68.68% | - |
| Mona monkey | 66.24% | 65.48% | 65.64% | - | 69.11% | 69.39% | 68.98% | - |
| De Brazza's monkey | 65.81% | 65.19% | 65.26% | - | 68.32% | 68.89% | 68.20% | - |
| Hanuman langur | 65.74% | 65.16% | 65.26% | - | 68.35% | 68.26% | 67.92% | - |
| Proboscis monkey | 54.38% | 52.97% | 53.76% | - | 56.79% | 56.20% | 55.88% | - |
| White-faced saki | 51.23% | 51.75% | 51.46% | - | 55.09% | 55.98% | 54.98% | - |
| Black-handed spider monkey | 50.56% | 50.82% | 50.46% | - | 54.75% | 54.98% | 54.18% | - |
| Tufted capuchin | 50.25% | 50.27% | 50.04% | - | 54.10% | 54.60% | 53.74% | - |
| Mantled howler monkey | 50.15% | 50.22% | 49.91% | - | 54.19% | 54.19% | 53.49% | - |
| White-fronted capuchin | 49.71% | 49.91% | 49.47% | - | 53.57% | 53.78% | 52.84% | - |
| Bolivian titi | 49.45% | 49.69% | 49.38% | - | 53.27% | 53.89% | 52.80% | - |
| Tamarin | 48.89% | 49.11% | 49.00% | - | 52.77% | 53.27% | 52.40% | - |

Supplementary Table S2: Top 30 most common species in which donor and acceptor splice sites are conserved in different datasets, including the "Random" baseline dataset (for comparison). To compute these species, we only used splice sites from GENCODE, RefSeq, CHESS conserved in less than 50 species that appear to have a random-like conservation pattern. Each entry shows the proportion of splice sites that have their canonical dinucleotides conserved in certain species relative to the total number of sites in the respective category. The first half of the table shows sites from protein-coding genes, the second one shows sites from lncRNAs. For the second part, the "Random" column show dashes since the same set of generated sites were used as a baseline for comparing against both coding and non-coding genes.

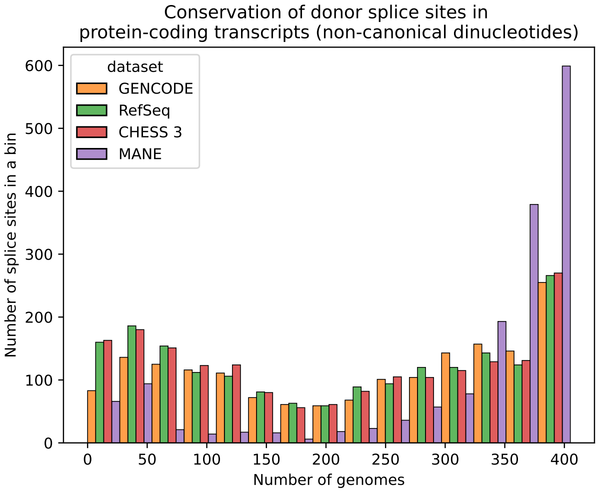


(a)

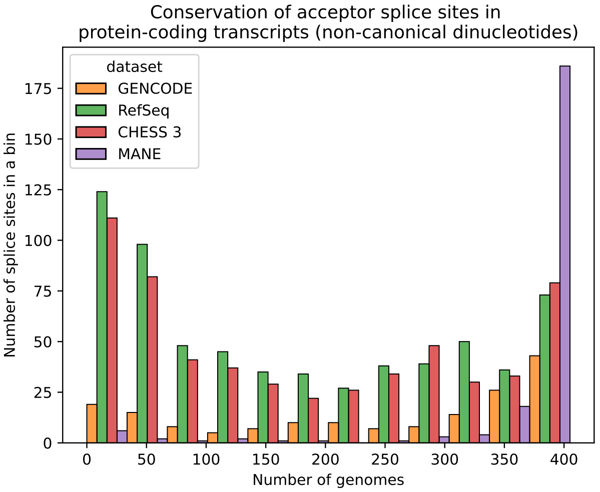


(b)

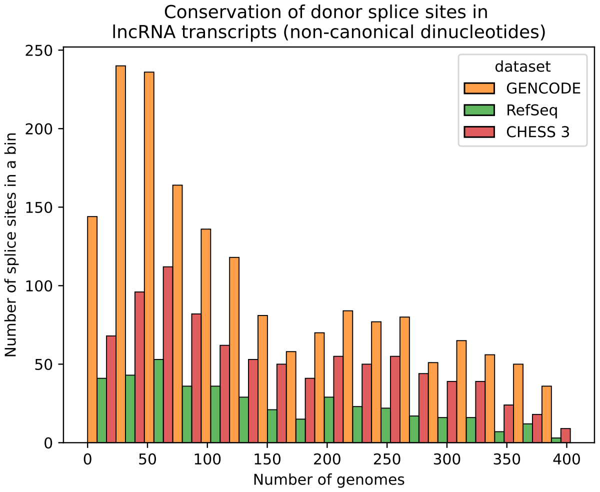


(c)

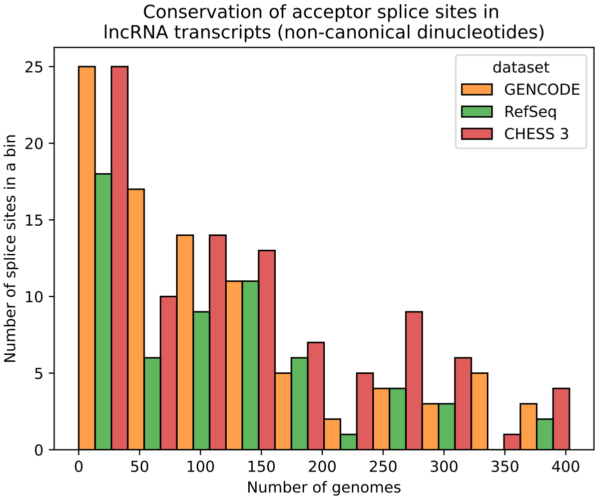


(d)

Supplementary Figure S1: The number of human splice sites with non-canonical dinucleotides in place of GT/AG conserved in 470 mammals, computed for donor (a) and acceptor (b) sites of protein-coding genes, and donor (c) and acceptor (d) sites of lncRNAs. Each point shows a number of splice sites conserved (y-axis) in a given number of species (x-axis), and data points are binned. For protein-coding genes, we created subsets GENCODE, RefSeq, and CHESS 3 from which we removed MANE annotations because each of these datasets is a superset of MANE; the resulting datasets are designated as GENCODE$^*$, RefSeq$^*$, and CHESS 3$^*$ correspondingly.

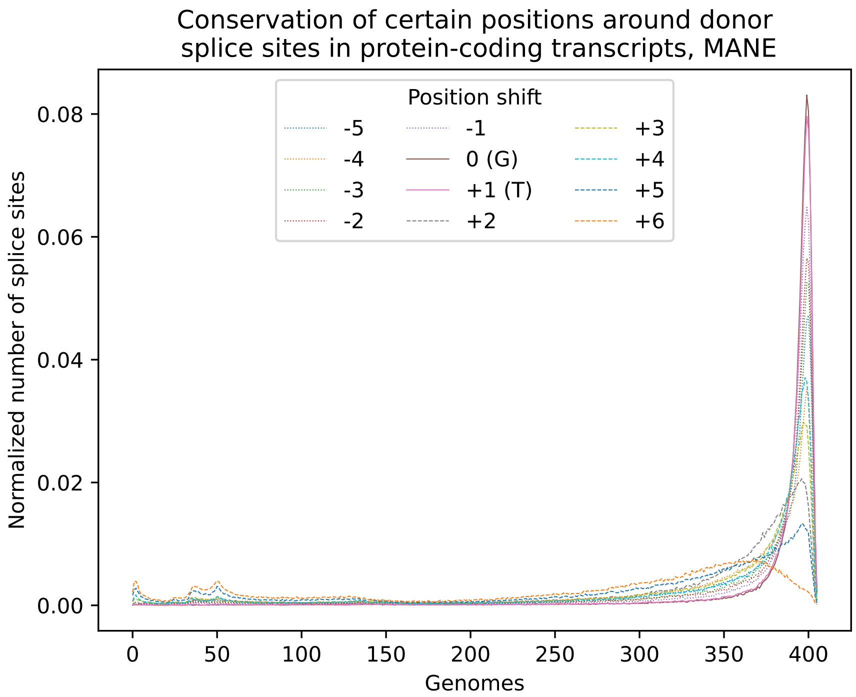


(a)

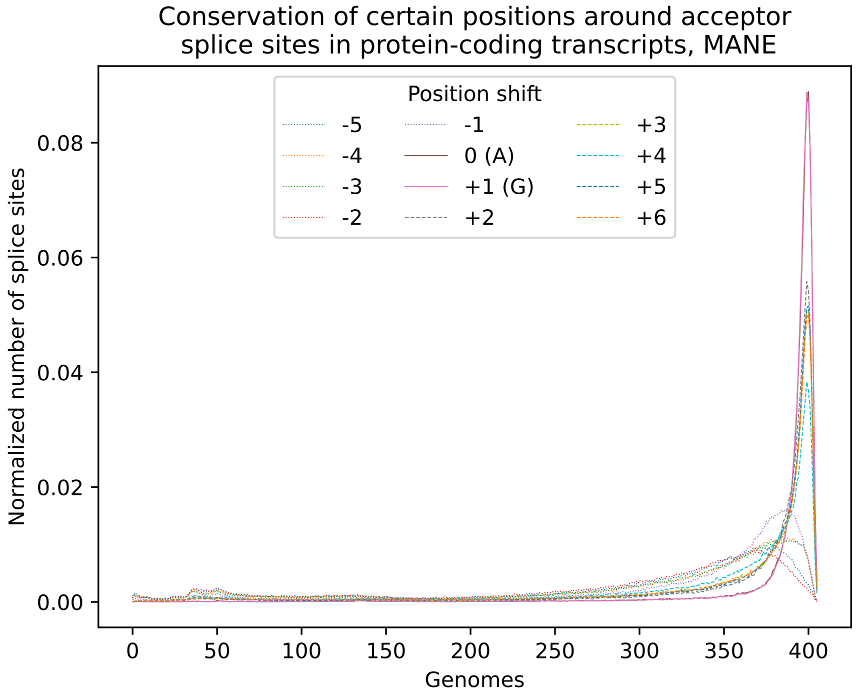


(b)

Supplementary Figure S2: Distribution of the number of donor (a) and acceptor (b) splice sites with a certain position around splice site motif conserved in a given number of species, for protein-coding genes from the MANE dataset. Each line represents the conservation of a position either down- or upstream of the "canonical" dinucleotides. For example, for donor splice sites 0 is usually "G", 1 is "T", and -1 is the first nucleotide upstream of the splice site. These nucleotides are shown in parentheses in the legends of the corresponding positions. Numbers are normalized by the total number of sites in the corresponding dataset in each category.

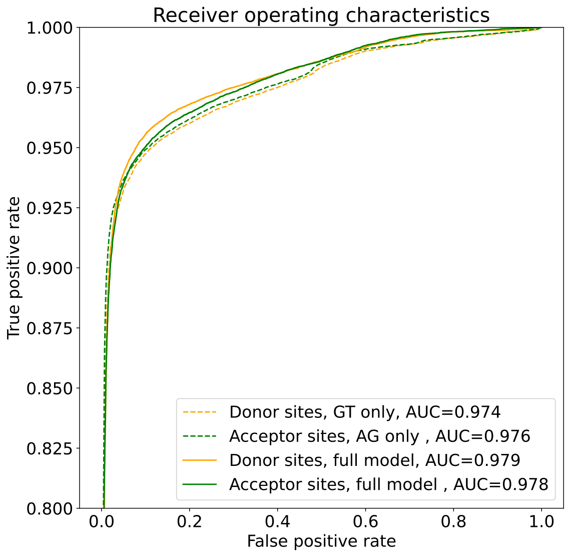


Supplementary Figure S3: Receiver operating characteristic (ROC) curve for logistic regression models trained to distinguish between GT/AG sites chosen at random from intronic sequences and splice sites from MANE. Dashed lines correspond to the models using only the conservation of the canonical dinucleotides themselves, while solid ones show the performance of the model taking into account the conservation of the positions in a larger window.

| Model | Recall | Precision | F-score |
|---|---|---|---|
| *Canonical dinucleotides only* | | | |
| Donor sites | 0.946 | 0.911 | 0.928 |
| Acceptor sites | 0.943 | 0.903 | 0.923 |
| *Full model* | | | |
| Donor sites | 0.943 | 0.923 | 0.933 |
| Acceptor sites | 0.949 | 0.949 | 0.949 |

Supplementary Table S3: Precision and recall of models classifying splice sites as either "well-supported" or "less-supported" on the test dataset containing. The first two rows show the performance of the models taking into account the canonical dinucleotides GT/AG only, while the next two rows correspond to the model considering a larger window. The classification threshold for probability is 0.5. Donor (acceptor) test set consisted of 36,000 false examples, randomly generated from intronic sequences, and 36,000 positive examples from MANE.

| Dataset | All splice sites | | "Well-supported" splice sites | |
|---|---|---|---|---|
| *Protein-Coding* | Donor | Acceptor | Donor | Acceptor |
| MANE | 1,617 | 225 | - | - |
| GENCODE* | 1,737 | 172 | 556 | 83 |
| RefSeq* | 1,877 | 647 | 571 | 157 |
| CHESS 3* | 1,874 | 572 | 563 | 149 |
| *lncRNA* | | | | |
| MANE | - | - | - | - |
| GENCODE | 1,746 | 89 | 169 | 8 |
| RefSeq | 419 | 60 | 28 | 3 |
| CHESS 3 | 897 | 94 | 66 | 9 |

Supplementary Table S4: Summary statistics of splice site conservation analysis, non-canonical splice sites. The second and third columns represent the total number of donor and acceptor sites in each dataset. The third and fourth columns show the number of donor and acceptor splice sites classified as "well-supported" by our model.

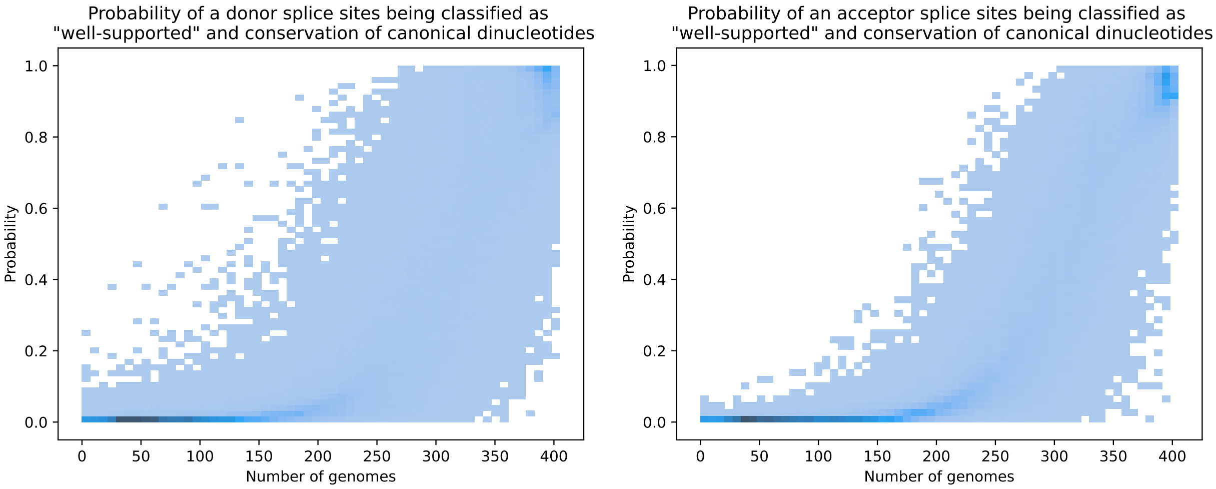


(a) (b)

Supplementary Figure S4: The relationship between the probability of a donor (a) and acceptor (b) splice sites classified as "well-supported" and the number of genomes in which the canonical dinucleotides are conserved. We used the "full model" that considers the conservation of the soliciting motif, as well as the canonical dinucleotides. Each panel depicts donor (acceptor) splice sites from protein-coding genes of GENCODE*, RefSeq*, and CHESS 3*, as well as form lncRNA genes of GENCODE, RefSeq, and CHESS 3.

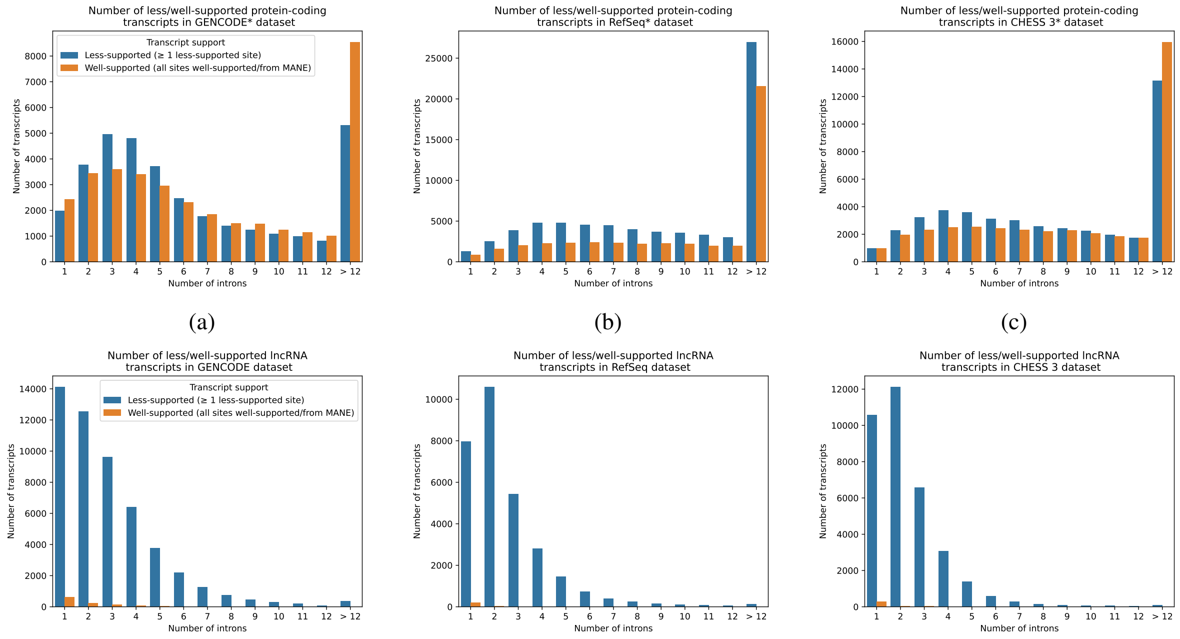


(d) (e) (f)

Supplementary Figure S5: The number of protein-coding and lncRNA transcripts in GENCODE, RefSeq, and CHESS 3 with a certain number of introns split by their support status. A transcript is defined as well-supported if all of its splice sites either are present in the MANE dataset or classified as "well-supported" by our model, otherwise the transcript is "less-supported. Panels (a)-(c) show transcripts from protein-coding genes, while (d)-(f) represent lncRNAs.

| Category | Donor sites | | | | Acceptor sites | | | |
|---|---|---|---|---|---|---|---|---|
| Inside a MANE exon | No | | Yes | | No | | Yes | |
| "Well-supported" | No | Yes | No | Yes | No | Yes | No | Yes |
| GENCODE* | 20,560 | 7,650 | 2,815 | 2,966 | 13,480 | 4,969 | 3,370 | 5,044 |
| RefSeq* | 32,813 | 24,988 | 2,510 | 2,549 | 25,315 | 21,741 | 2,859 | 4,245 |
| CHESS 3* | 23,958 | 21,170 | 2,667 | 2,679 | 19,566 | 18,330 | 2,988 | 4,575 |

Supplementary Table S5: Summary statistics of splice site conservation analysis, with sites being split by whether they are located inside a MANE exon or an intron. Each column shows either the number of "well-supported or "less-supported" splice sites from each dataset, located inside or outside MANE exons, in each dataset in both categories (donor and acceptor).

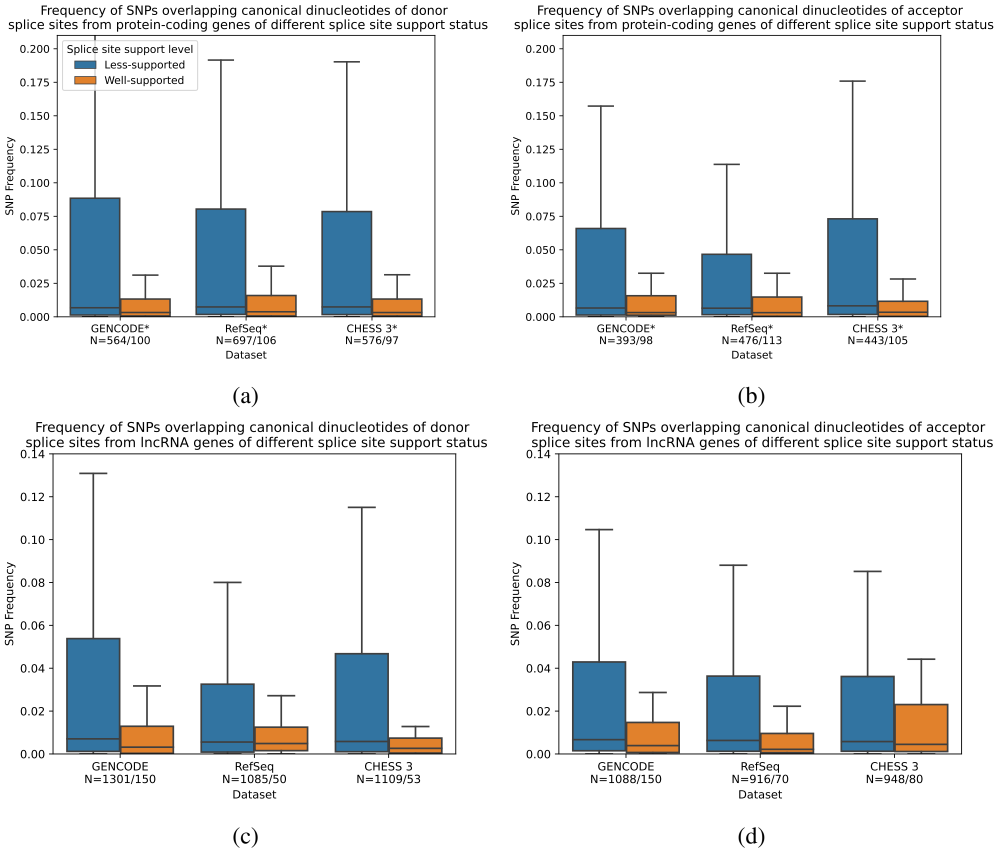


Supplementary Figure S6: Box plots showing the distribution of SNP frequencies from the gnomAD dataset overlapping the canonical dinucleotides GT/AG of splice sites of different support status as determined by our model. Panel (a) and (b) show donor and acceptor sites from protein-coding genes, while (c) and (d) illustrate donor and acceptor sites from lncRNAs. Each pair of boxplots shows the distribution of frequencies of splice sites from a particular dataset and support categories; numbers before the dataset title show the number of SNPs overlapping splice sites in well-supported/less-supported categories. Each box plot shows the median (solid horizontal line), the interquartile range (solid top and bottom borders of the box), and maximum values within 95% percentile (whiskers), outliers are not shown.

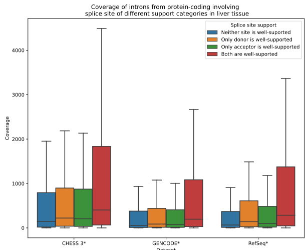


(a)

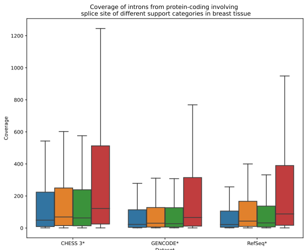


(b)

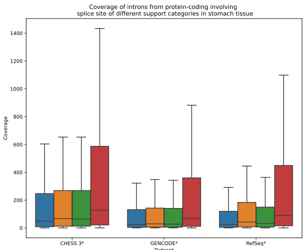


(c)

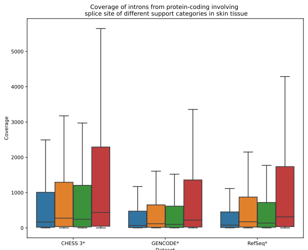


(d)

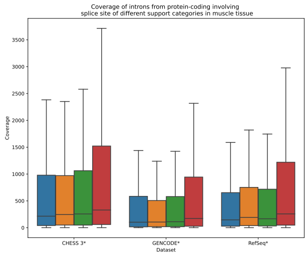


(e)

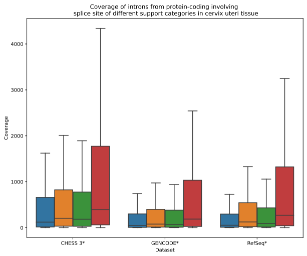


(f)

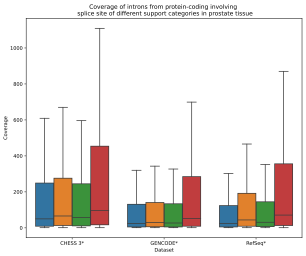


(g)

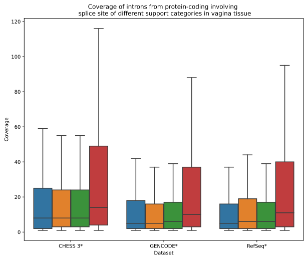


(h)

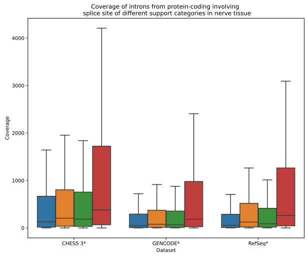


(i)

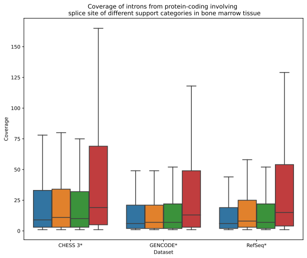


(j)

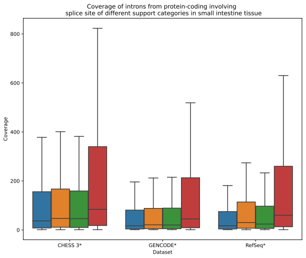


(k)

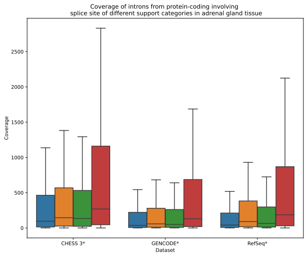


(l)

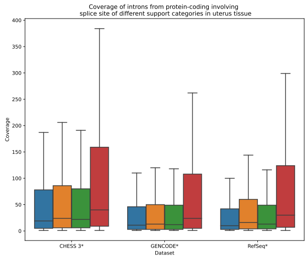


(m)

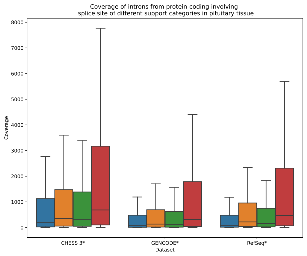


(n)

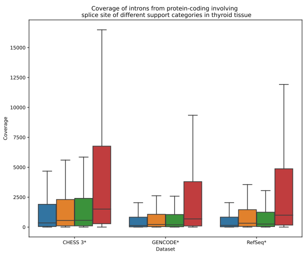


(o)

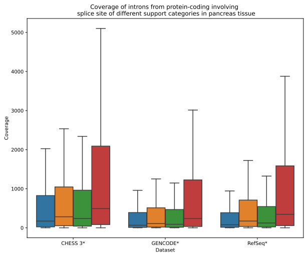


(p)

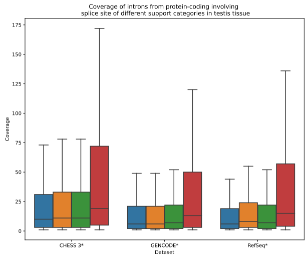


(q)

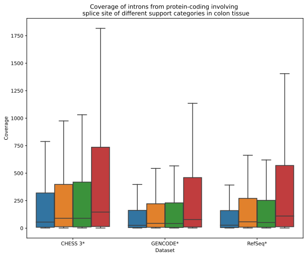


(r)

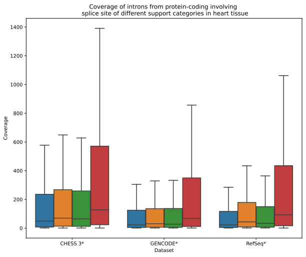


(s)

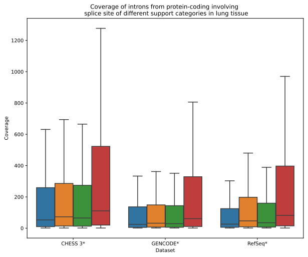


(t)

Supplementary Figure S7: Box plots showing coverage of introns from protein-coding genes of GENCODE*, RefSeq*, from CHESS 3* across different tissues. Each box plot shows the median (solid horizontal line), the interquartile range (solid top and bottom borders of the box), and minimum and maximum values within ±1.5 of the interquartile range (whiskers); outliers are not shown.

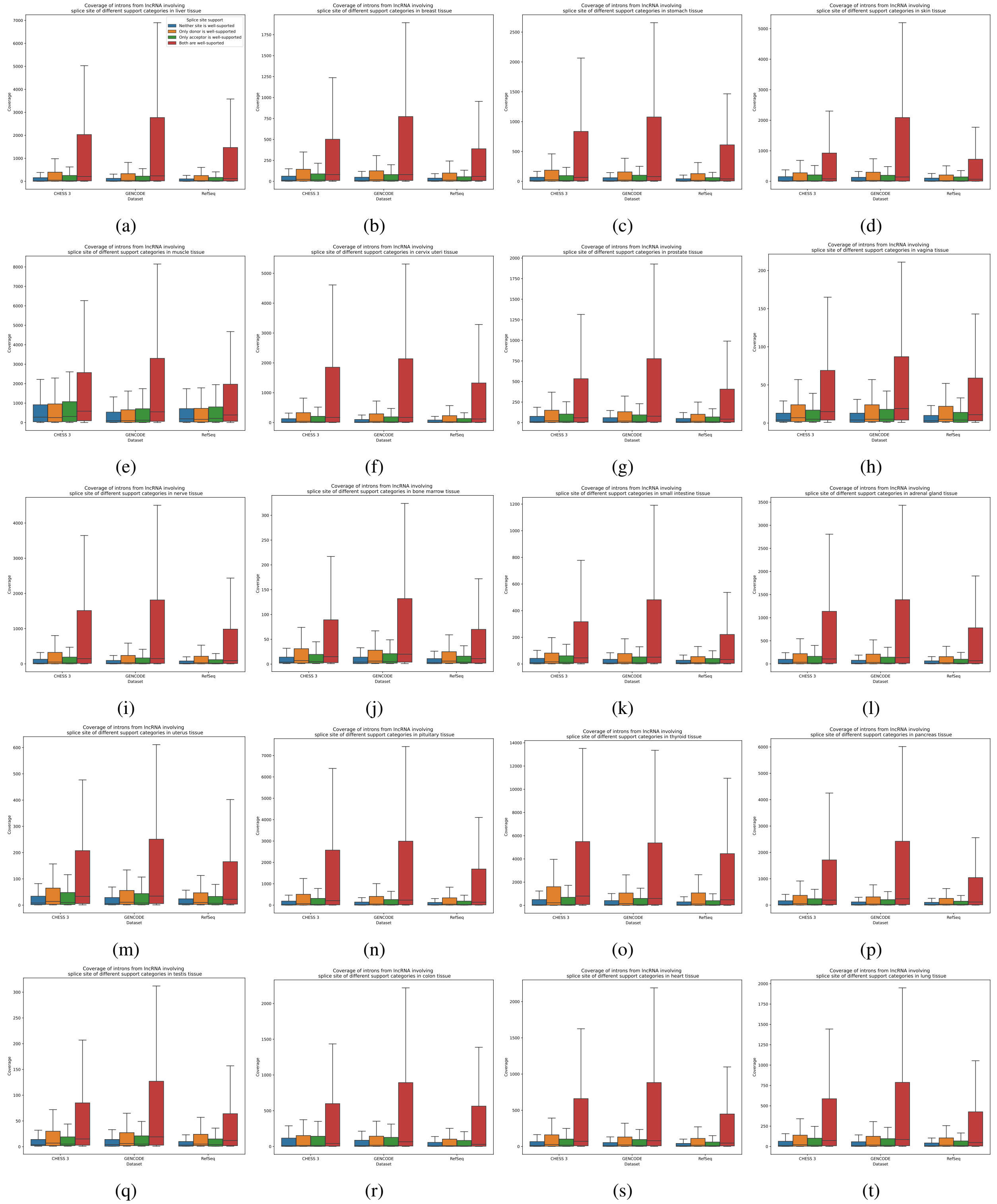


Supplementary Figure S8: Box plots showing coverage of introns from lncRNAs of GENCODE*, RefSeq*, from CHESS 3* across different tissues. Each box plot shows the median (solid horizontal line), the interquartile range (solid top and bottom borders of the box), and minimum and maximum values within ±1.5 of the interquartile range (whiskers); outliers are not shown.

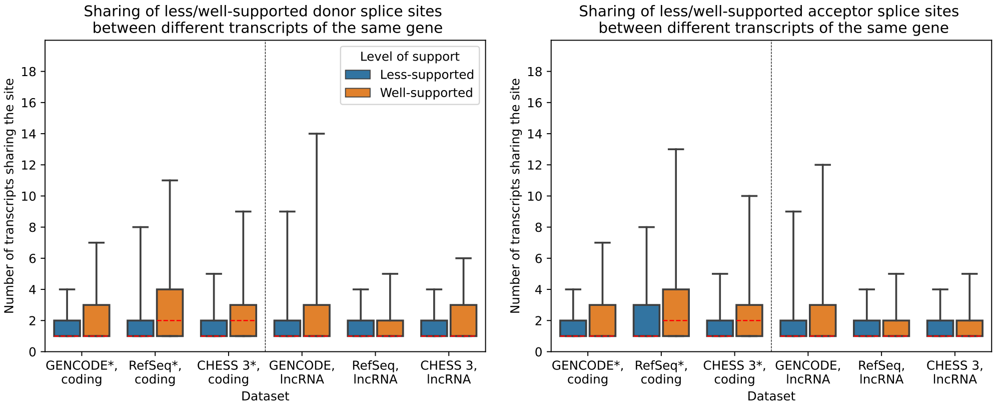


(a) (b)

Supplementary Figure S9: Box plots showing the distribution of the number of transcripts sharing a certain donor (a) and acceptor (b) splice site, in GENCODE*, RefSeq*, and CHESS 3* datasets. The left part of each panel shows splice sites from protein-coding genes, while the right represents lncRNAs. Each box plot shows the median (dashed red line), the interquartile range (solid top and bottom borders of the box), and maximum values within 95% percentile (whiskers), outliers are not shown.

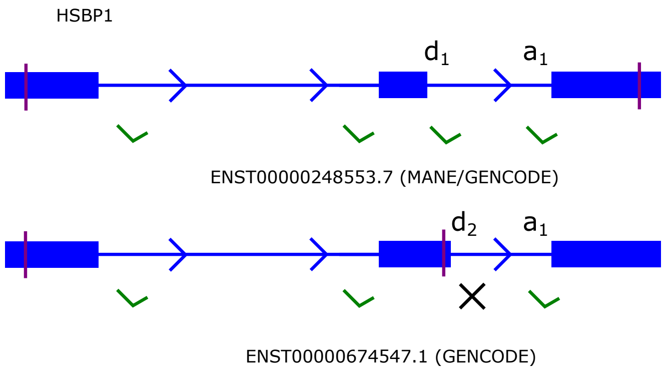


(a)

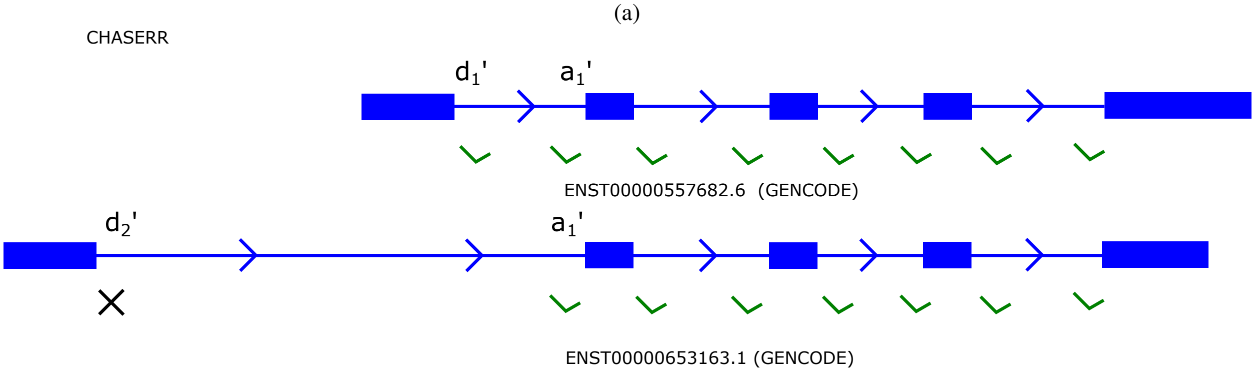


(b)

Supplementary Figure S10: This figure shows the structure of the transcripts containing less-supported splice sites that we believe could be misannotated. Solid blue boxes show exons, and blue arrows represent introns; green ticks indicate "well-supported" splice sites, while black crosses indicate “less-supported” ones, and vertical purple lines indicate start/stop codons. Figures are not drawn to an exact scale. Panel (a) shows two isoforms of the heat shock protein family B (small) member 1, or HSPB1 gene: MANE, ID ENST00000248553.7, upper transcript, and its alternative isoform from GENCODE, ID ENST00000674547.1, lower transcript that differs by a location of a donor splice site $d_2$. This alternative site results in a premature stop codon terminating the protein. The resulting protein is only 143 amino acids long. compared to the MANE isoform which contains 203 amino acids. Panel (b) shows two isoforms of lncRNA CHASERR from GENCODE: the upper isoform is ENST00000653163.1 and all its splice sites are “well-supported” according to our model. The lower isoform ENST00000557682.6 differs from the former transcript by the location of its first donor splice site ($d'_2$), which is less-supported. This isoform also occupies a genetic locus that is 4.7 times longer than its well-supported counterpart.

| | Gene | HSBP1 | | CHASERR | |
|---|---|---|---|---|---|
| | Intron | $d_1$ (✓) $a_1$ (✓) | $d_2$(×) $a_1$ (✓) | $d'_1$ (✓) $a'_1$ (✓) | $d'_2$ (×) $a'_1$ (✓) |
| Tissue | Spleen | 14,477,176 | 17 | 89,208 | - |
| | Salivary Gland | 13,461,418 | 5 | 123,743 | - |
| | Blood | 8,234,251 | 11 | 109,608 | - |
| | Blood Vessel | 5,605,781 | 12 | 32,065 | - |
| | Ovary | 5,154,826 | 8 | 29,174 | - |
| | Thyroid | 3,200,006 | 2 | 166,257 | 2 |
| | Nerve | 3,080,882 | 15 | 41,658 | - |
| | Pituitary | 2,829,673 | - | 79,553 | - |
| | Breast | 1,832,637 | - | 12,905 | - |
| | Pancreas | 1,576,185 | - | 54,498 | - |
| | Liver | 1,538,312 | 24 | 53,020 | - |
| | Heart | 1,214,850 | - | 11,977 | - |
| | Skin | 908,380 | - | 51,320 | - |
| | Cervix Uteri | 876,715 | - | 53,241 | - |
| | Fallopian Tube | 852,013 | - | 11,997 | - |
| | Bladder | 763,279 | 1 | 15,228 | - |
| | Adrenal Gland | 743,273 | 1 | 36,980 | - |
| | Brain | 680,486 | - | 14,915 | - |
| | Lung | 521,222 | - | 10,311 | - |
| | Small Intestine | 465,708 | - | 7,964 | - |
| | Stomach | 404,787 | - | 15,357 | - |
| | Esophagus | 362,020 | - | 9,530 | - |
| | Kidney | 309,843 | - | 13,341 | - |
| | Adipose Tissue | 303,877 | - | 5,161 | - |
| | Colon | 288,139 | - | 12,308 | - |
| | Muscle | 211,252 | - | 22,579 | - |
| | Uterus | 182,724 | - | 3,794 | - |
| | Prostate | 175,402 | - | 10,614 | - |
| | Bone Marrow | 90,179 | - | 1,011 | - |
| | Testis | 80,903 | - | 969 | - |
| | Vagina | 40,997 | - | 747 | - |

Supplementary Table S6: This table shows the aggregated number of reads across all samples in each tissue in the GTEx dataset supporting the exon junction described in the subsection "Case study: spotting potentially suspicious isoforms" of the main text. Columns 2 and 3 show the number of reads supporting introns ($d_1$, $a_1$) and ($d_2$, $a_1$) of the gene HSPB1, both sharing the same acceptor splice site; however, the donor site $d_1$ is well-supported, while the donor splice of $d_2$ is less-supported. Ticks indicate well-supported splice sites, while crosses show less-supported sites. Columns 4 and 5 depict the same date but for lncRNA gene CHASERR, and the support status is analogous: the donor site $d'_1$ is well-supported, while the donor splice of $d'_2$ is less-supported Supplementary Figure S10 shows the structure of these two genes and the locations of the splice sites inside each respective gene.
